# Endocytosis in *Trypanosoma cruzi* Depends on Proper Recruitment and Regulation of Functionally Redundant Myosin Motors

**DOI:** 10.1101/2022.11.17.517012

**Authors:** Nathan M. Chasen, Menna G. Etheridge, Paul C. Campbell, Christopher L. de Graffenried, Kingsley Bimpeh, Kelly M. Hines, Ronald D. Etheridge

**Author notes:** **<u>Email:</u>** Nathan M. Chasen, Menna G. Etheridge, Kingsley Bimpeh, Kelly M. Hines, Paul C. Campbell, Christopher L. de Graffenried.

## Abstract

Utilized by the free-living kinetoplastid *Bodo saltans* to feed on bacterial prey, the cyto<u>s</u>tome-cyto<u>p</u>harynx <u>c</u>omplex (SPC) is an endocytic organelle absent from all human trypanosomatid pathogens save *Trypanosoma cruzi.* Building upon our previous work identifying the myosin motor MyoF as the first enzymatic component of the *T. cruzi* SPC, we sought to expand our understanding of this distinct organelle by identifying additional protein machinery which contribute to the endocytic process. While deletion of MyoF alone did not fully ablate endocytosis, we found that deletion of both MyoF and the similarly localized MyoC produced an endocytic-null phenotype that was rescued upon complementation. To identify potential regulatory components of this motor complex, we pulled down MyoF and identified an SPC-targeted protein that contained an annotated EF-hand calcium-binding motif that was conserved across a wide range of protozoan lineages. Surprisingly, deletion of this <u>m</u>yosin <u>a</u>ssociated <u>p</u>rotein (MyAP) alone was sufficient to produce an endocytic-null phenotype, which we were able to fully rescue via complementation. The deletion of MyAP also caused the mis-localization of both cytopharynx myosins to the cytosol. While MyAP lacking the EF-hand domain was unable to complement endocytosis, it was sufficient to restore proper myosin localization. This suggested that MyAP plays two distinct roles, one in targeting myosins to the SPC and a second in regulating myosin motor activity. Transmission electron microscopy also revealed that endocytic-null mutants lacked the electron lucent lipid inclusions typically seen in the pre-lysosomal reservosomes of *T. cruzi* epimastigotes. Mass spectrometry based lipidomic analysis subsequently revealed a dramatic reduction in the scavenged cholesterol content in the endocytic-null mutants, which can be attributed to an inability to endocytose exogenous lipid-protein complexes for storage in the reservosomes. Overall, this work showcases the first viable endocytic-null mutants generated in *T. cruzi* through specific gene deletion and highlights the feasibility of leveraging this strategy towards a full dissection of the endocytic machinery and biogenesis of the SPC.

**Importance:** *Trypanosoma cruzi* chronically infects over 7 million people in the Americas and current therapeutics are insufficient to effectively cure infection. The lack of progress in developing effective vaccines or drug treatments is due, in part, to longstanding technical limitations in studying this parasite and a lack of resources committed to support research and eradication efforts. As part of its parasitic lifestyle, *T. cruzi* is forced to obtain basic nutrients directly from its host environment, making the development of methods to block nutrient uptake an attractive strategy to control parasite growth and transmission. While the bulk uptake of complex nutrients by *T. cruzi* occurs via an endocytic structure, often referred to as the cyto<u>s</u>tome-cyto<u>p</u>harynx <u>c</u>omplex (SPC), how exactly this tubular endocytic organelle functions at a mechanistic level has remained a mystery. In this work, we investigated the contribution of several SPC targeted myosin motors and an associated protein factor to endocytic activity. By identifying and characterizing the molecular machinery responsible for nutrient uptake, we hope to both expand our basic understanding of how this deadly pathogen acquires essential nutrients from its host, while also revealing new potential therapeutic targets to impede nutrient uptake.

## Introduction

*Trypanosoma cruzi*, the etiological agent of Chagas disease, is a protozoan parasite that chronically infects upwards of 7 million people in the Americas resulting in an estimated 50,000 deaths annually (1, 2). Unlike heavily studied salivarian trypanosomatids such as *Trypanosoma brucei* and *Leishmania spp.* which invade their human hosts directly through the proboscis of their hematophagous insect vectors, the stercorarian *Trypanosoma cruzi* is released in the feces of its blood feeding reduviid vector and must contaminate the bite wound or nearby mucosal membranes to infect its vertebrate host (3, 4). Over decades, chronic parasite infection can lead to the destruction of smooth and cardiac muscle tissue, ultimately manifesting as various mega viscera or cardiac disease in approximately 30% of those infected (5). Unfortunately, no effective vaccines are available to prevent infection and the chemical therapeutics currently in use are often toxic and ineffective against chronic infection (6, 7). Recent work has also highlighted parasite dormancy in the mammalian host as a potential mechanism by which *T. cruzi* may be able to resist clearance by the few chemotherapeutic options currently available, further supporting the need for a better understanding of this parasite’s basic biology (8).

One intriguing dimension of *T. cruzi’s* fundamental physiology which remains poorly understood is the mechanism by which the parasite acquires nutrients from it host environment. Unlike its salivarian cousins (*Trypanosoma brucei* and *Leishmania spp.*) which have repurposed their flagellar pocket membrane to be the sole location for endocytosis and exocytosis (9–11), *T. cruzi* utilizes an ancestral form of phagocytosis operating via a flagellar pocket adjacent organelle known as the cyto<u>s</u>tome-cyto<u>p</u>harynx com<u>p</u>lex (SPC) which is still used by its free-living relatives (e.g. *Bodo saltans*) to capture and consume bacterial prey (reviewed in (12)). The SPC begins as an opening on the parasite surface (cytostome) and is followed by a dynamic tubular membrane invagination (cytopharynx) through which captured and endocytosed material is brought into the cell and ultimately digested (13). Connecting the opening of the flagellar pocket to the cytostome entrance is a unique cholesterol and glycan rich plasma membrane subdomain known as the pre-oral ridge (POR) that is compositionally distinct from the rest of the parasite surface membrane and originates at the base of the flagellar pocket via vesicular fusion. It is on this POR at the parasite surface that complex nutrients are thought to be captured by, as yet unknown, membrane receptors prior to being drawn down into the SPC. The endocytic complex ultimately terminates with the budding of vesicles that are trafficked to the pre-lysosomal storage structures known as reservosomes to await digestion (14, 15). Undergirding this phagocytic structure are two sets of microtubule root fibers; one known as the cytostomal quartet (CyQ) which begins at the basal body, winds up the flagellar pocket and runs beneath the POR membrane before descending into the parasite body while the second rootlet known as the cytostomal triplet (CyT) originates adjacent to the cytostome opening itself and tracks alongside the CyQ forming a “gutter” within which lies the cytopharynx membrane tubule (16). Among single celled protozoans, these microtubule rootlets often facilitate the proper positioning and function of various subcellular organelles (reviewed in (17)). Up until recently, the vast majority of our understanding of the SPC apparatus was gleaned from structural examinations using electron microscopy-based techniques (12). These methods, however, were unable to provide mechanistic insight into how this organelle is able to capture and pull in endocytosed material. Our group has previously published both the identification of the first known SPC targeted proteins (18) as well as a follow-up characterization of the first enzymatic component, a myosin motor known as MyoF, which contributes to SPC mediated endocytosis (19). In this prior work, we also overexpressed an enzymatically dead rigor-mutant of MyoF and found that it completely blocked measurable endocytic activity. Counterintuitively, however, parasites in which MyoF was directly knocked out (KO) still demonstrated measurable, yet highly diminished, endocytosis suggesting that functionally redundant myosin motors may be contributing to this activity and that the rigor-mutant was acting in a dominant-negative fashion. As a result, we have identified three additional SPC targeted myosin motors and demonstrated that two of these are positioned at the pre-oral ridge (MyoB and MyoE), while a third myosin isoform (MyoC) localizes to the cytopharynx.

In this study, we have continued our analysis of the cytopharynx targeted MyoF and MyoC by generating a double deletion mutant in *T. cruzi.* Parasites lacking both MyoF and MyoC were completely devoid of measurable endocytic activity that was subsequently restored upon individual gene complementation. To further characterize the molecular complexes regulating these motors, we performed co-immunoprecipitation (co-IP) of MyoF and identified an SPC cytopharynx targeted <u>m</u>yosin <u>a</u>ssociated <u>p</u>rotein (MyAP). Unlike most SPC components identified to date, orthologs of MyAP were identified in a variety of distantly related protozoans suggesting a potentially conserved or ancestral role in protozoal phagotrophy. Deletion of MyAP in epimastigotes gave rise to an endocytic-null phenotype that mirrored the double deletion MyoF and MyoC KO mutants. We utilized both super-resolution and expansion microscopy to demonstrate that MyAP and the myosin motors are targeted specifically to the SPC microtubule rootlets and that loss of MyAP resulted in parasites no longer being able to properly target MyoF and MyoC to these microtubules. While a loss of endocytosis did not directly impact parasite growth *in vitro*, it led to a dramatic change in the apparent lipid composition of the pre-lysosomal reservosomes observed in transmission electron-microscopy (TEM) images. High-resolution mass spectrometry (MS) confirmed this observation and revealed a striking decrease in scavenged host cholesterol in endocytic-null mutants, whereas the levels of endogenously synthesized ergosterol (20) remained unchanged. Broadly, this work demonstrates the first use of gene deletion to produce endocytic-null mutants in *T. cruzi* and lays the groundwork for a full dissection of the SPC, including the molecular components essential for both the biogenesis and function of this enigmatic endocytic organelle.

## Results

### The cytopharynx targeted myosins MyoF and MyoC are necessary for endocytic function

In our prior report on the functional characterization of the cytopharynx targeted MyoF in *Trypanosoma cruzi*, we over-expressed a catalytically dead rigor-mutant of MyoF that had the effect of completely blocking measurable endocytosis (19). In addition to the surprising revelation that this endocytic-null mutant exhibited no significant changes in growth or viability in culture, this observation led us to believe that MyoF was an essential component of the endocytic complex. However, after the generation of a homozygous knockout (KO) of MyoF, we still observed detectable levels of endocytosis, suggesting that redundant or compensatory myosin motor activity may be present. With this in mind, we localized the remaining orphan myosins in *T. cruzi* and observed that isoforms MyoB and MyoE localized to the <u>p</u>re-<u>o</u>ral <u>r</u>idge (POR) region of the cyto<u>s</u>tome-cyto<u>p</u>harynx <u>c</u>omplex (SPC) while MyoC targeted to the cytopharynx much like MyoF (**Figure 1** structural schematic and microscopy localization). This result bolstered our hypothesis that additional myosins were potentially involved in endocytosis and, as a test, we used CRISPR/Cas9 (**Figure 2A** methodology) to delete MyoC both alone (*ΔMyoC*) and in combination with MyoF (*ΔFΔC*) (**Figure 2B** and **2C** polymerase chain reaction (PCR) validation for MyoC and MyoF loci changes). We also complemented MyoC back into the double *ΔFΔC* background (*ΔFΔC::C-Ty*) and validated its proper localization to the cytopharynx using immunofluorescence microscopy (IFA) (**Figure 2D**). While the loss of MyoC alone had a negligible effect on the overall rate of endocytosis as compared to *ΔMyoF* (**Figure 2E** and quantified in **2G** (green *ΔMyoC* and blue *ΔMyoF*)), deletion of both MyoF and MyoC (*ΔFΔC*) resulted in a severe defect in endocytic rate on a par with our chemical inhibitor of endocytosis: the actin polymerization inhibitor cytochalasin D (+CytD) (**Figure 2F** and quantified in **2G** (gray: +CytD and red: *ΔFΔC*)). As expected, complementation of MyoC restored the endocytic rate to the original *ΔMyoF* levels (**Figure 2F** and quantified in **2G** (yellow)). As observed previously when we overexpressed the rigor-mutant of MyoF, loss of these motors and endocytic function as a whole, had no significant effect on parasite growth *in vitro* (**Supplementary Figure S1A** and **S1B**). Endocytosis, therefore, appears to rely primarily on MyoF, with MyoC also contributing significantly to this activity as evidenced by the double deletion mutant.

**Figure 1.**
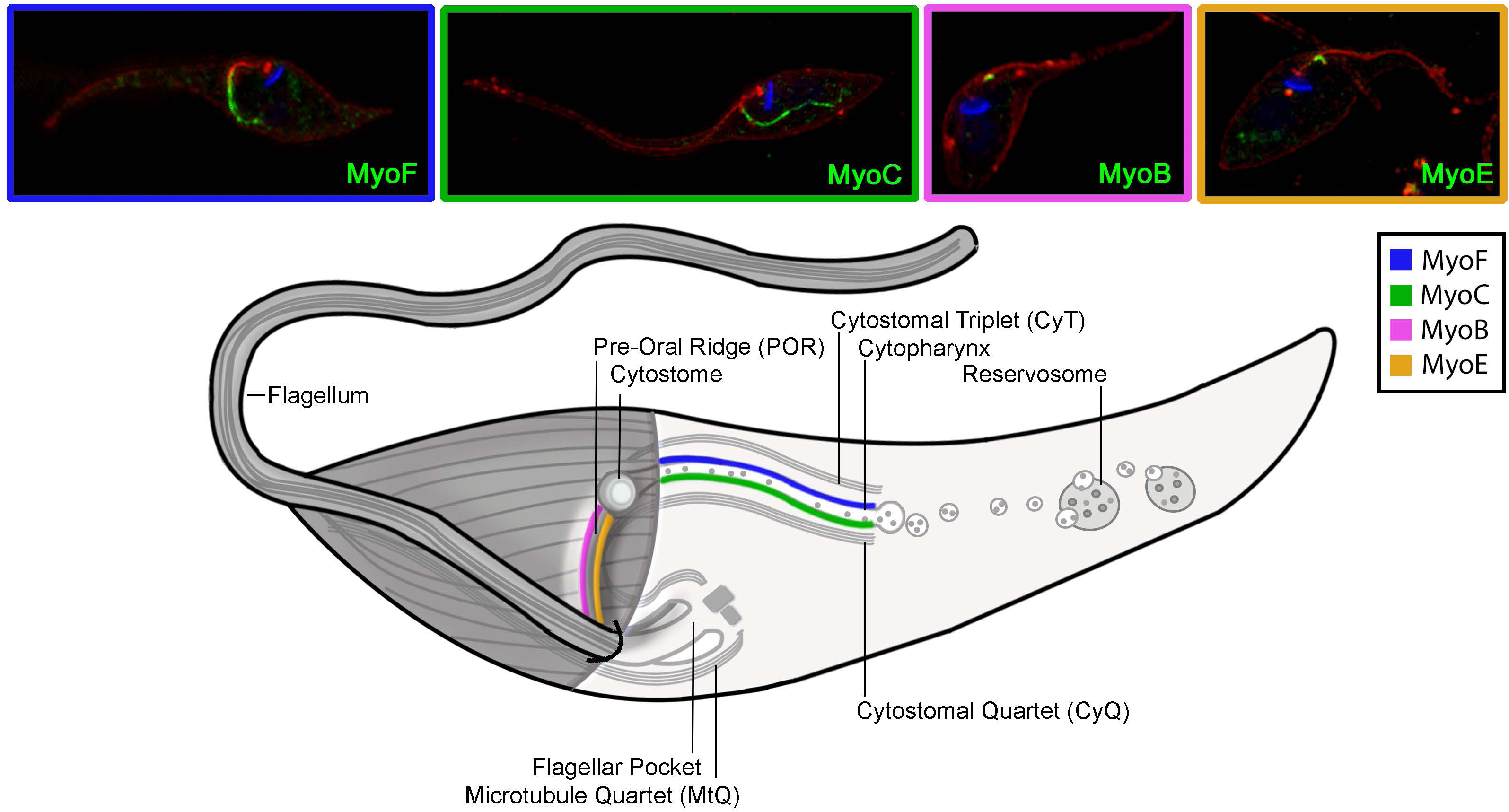
Schematic Summarizing the *Trypanosoma cruzi* SPC and Associated Myosins. Four myosins are associated with the cyto<u>s</u>tome cyto<u>p</u>harynx <u>c</u>omplex (SPC). MyoF (*blue*) and MyoC (*green*) localize to the cytopharynx portion of the SPC, while MyoB (*magenta*) and MyoE (*orange*) localize to the pre-oral ridge.

**Figure 2.**
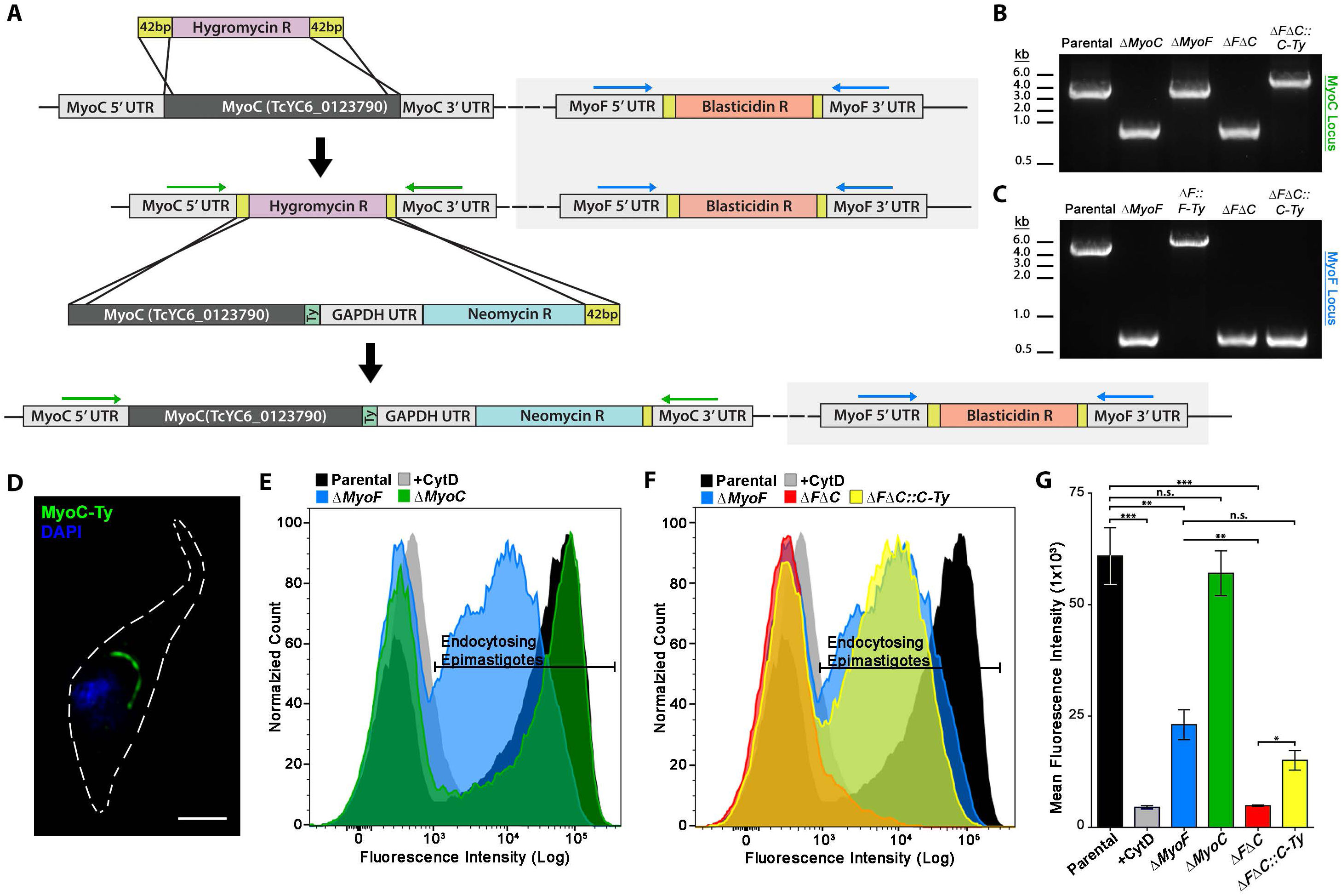
Double Knockouts for MyoF and MyoC Show a Synergistic Loss of SPC Endocytosis. **A.** Scheme for CRISPR/Cas9 gene deletion and complementation of MyoC in both Y-Strain and *ΔMyoF (gray tint*) backgrounds. Location of primers used for PCR verification of loci modifications are annotated (*green and blue arrows*). **B.** PCR amplification with MyoC screening primers (*green arrows in A*) shows replacement of both Parental loci (high-molecular-weight (MW) band) with the blasticidin resistance gene (low-MW band) in *ΔMyoC* and *ΔMyoFΔMyoC (ΔFΔC*) mutants. The higher MW PCR product amplified from the *ΔMyoFΔMyoC::MyoC-Ty (ΔFΔC::C-Ty*) mutants, shows correct insertion into the original *ΔMyoC* locus. **C.** PCR amplification with MyoF screening primers (*blue arrows in A*) shows replacement of the MyoF loci (high-MW band) with the blasticidin resistance cassette (low-MW band) in both the single and double knockout mutants. **D.** IFA of MyoC-Ty complemented epimastigotes shows localization to the distinct linear structure of the SPC. **E.** *ΔMyoC* epimastigotes (*green*) showed no significant reduction in SPC endocytic rate, whereas *ΔMyoF* (*blue*) epimastigotes show an endocytic rate reduction as previously described. Treatment with the actin polymerization inhibitor Cytochalasin D during the assay is used as a negative control for feeding assays throughout this manuscript (*grey*). **F.** *ΔFΔC* mutants (*red*) show a loss of detectable endocytosis, in contrast with the partial endocytic rate reduction seen in the *ΔMyoF (blue*) background. Complementation with MyoC-Ty in *ΔFΔC::C-Ty* mutants (*yellow*) resulted in a similar endocytic rate to that seen in the *ΔMyoF* single knockout background (*blue*). **G.** Quantification of endocytic rate in mutants shown in panels E and F show the synergy between MyoF and MyoC deletion in reducing SPC endocytosis. Scale bar 2 μm.

### A MyoF associated protein localizes to the microtubule rootlets of the SPC

To begin identifying the protein components associated with the MyoF motor complex, we carried out co-immunoprecipitation (co-IP) and mass spectrometry (MS) analysis of *T. cruzi* parasites overexpressing MyoF fused to the fluorescent protein mNeon and the Ty-tag epitope (**Figure 3A** top: protein gel and bottom: MS results). From this work, we identified a <u>m</u>yosin <u>a</u>ssociated <u>p</u>rotein (MyAP) which contains a putative calcium responsive paired EF-hand (EFh) domain (21) and a highly ubiquitous protein interaction module known as a leucine-rich repeat (LRR) (22) (**Figure 3B** top: MyAP AlphaFold prediction, EFh in blue and LRR in green and bottom: linear protein schematic). The presence of the EFh pair suggested a potential role in regulating motor function as these calmodulin-like domains are known to bind IQ motifs which are also predicted to be present in MyoF (23). In examining the genome of Y-strain *T. cruzi* (DTU II) (24, 25), we discovered the presence of two paralogs of MyAP (***a***: TcYC6_0120270 and ***b***: TcYC6_0120590) which are distinguished solely by the insertion of six amino acids (QYSSTQ) in the N-terminal portion of the subtype ***b*** protein (**Figure 3B** bottom: vertical red line denotes insertion location). We localized the overexpressed fusion of MyAP-mNeon-Ty (subtype ***b***) in transfected parasites and observed its targeting to the now-familiar SPC-like linear structure (**Figure 3C** green). To confirm co-localization of MyAP and components of the SPC, we first generated a mouse polyclonal antibody to a predicted antigenic region of MyAP (**Figure 3B** bottom; horizontal pink line denotes antigen). Using the resulting antibody, we conducted an IFA on parasites expressing the MyoF-mNeon-Ty fusion protein and observed clear overlap of the MyAP (red) and MyoF (green) fluorescent signals (**Figure 3D**). In order to localize MyAP with greater precision within the SPC itself, we conducted expansion microscopy on *T. cruzi* epimastigotes combined with an IFA against the Ty-epitope using our MyAP-Ty complemented line. Utilizing the TAT-1 anti-α-tubulin antibody (26, 27) to highlight parasite microtubules, we were able to demonstrate that MyAP-Ty specifically targets to the microtubule rootlets rather than the membrane tubule of the SPC (**Figure 3E** left panel). In addition, we examined the localization of both MyoF-Ty and MyoC-Ty using this same methodology and found that these motors also associated with the SPC microtubules (**Figure 3E** middle and right panels respectively) thus validating previous observations of MyoF using immuno-electron microscopy (28). Additional representative images demonstrating MyAP, MyoF and MyoC localizations are also presented in **Supplementary Figures S2A, S2B** and **S2C** respectively. As a negative control, the untagged parental and *ΔMyAP* strains failed to show any appreciable fluorescent signal on the cytostomal microtubules (**Supplementary Figures S2D** and **S2E** respectively). To further demonstrate MyAP’s association with the parasite microtubular cytoskeleton, we fractionated *T. cruzi* epimastigotes into detergent soluble and insoluble cytoskeletal fractions and, using Western blot analysis, observed an enrichment of MyAP with the cytoskeletal fraction with the protease Cruzipain serving as a marker for the detergent soluble fraction (**Supplementary Figure S3**) (29).

**Figure 3.**
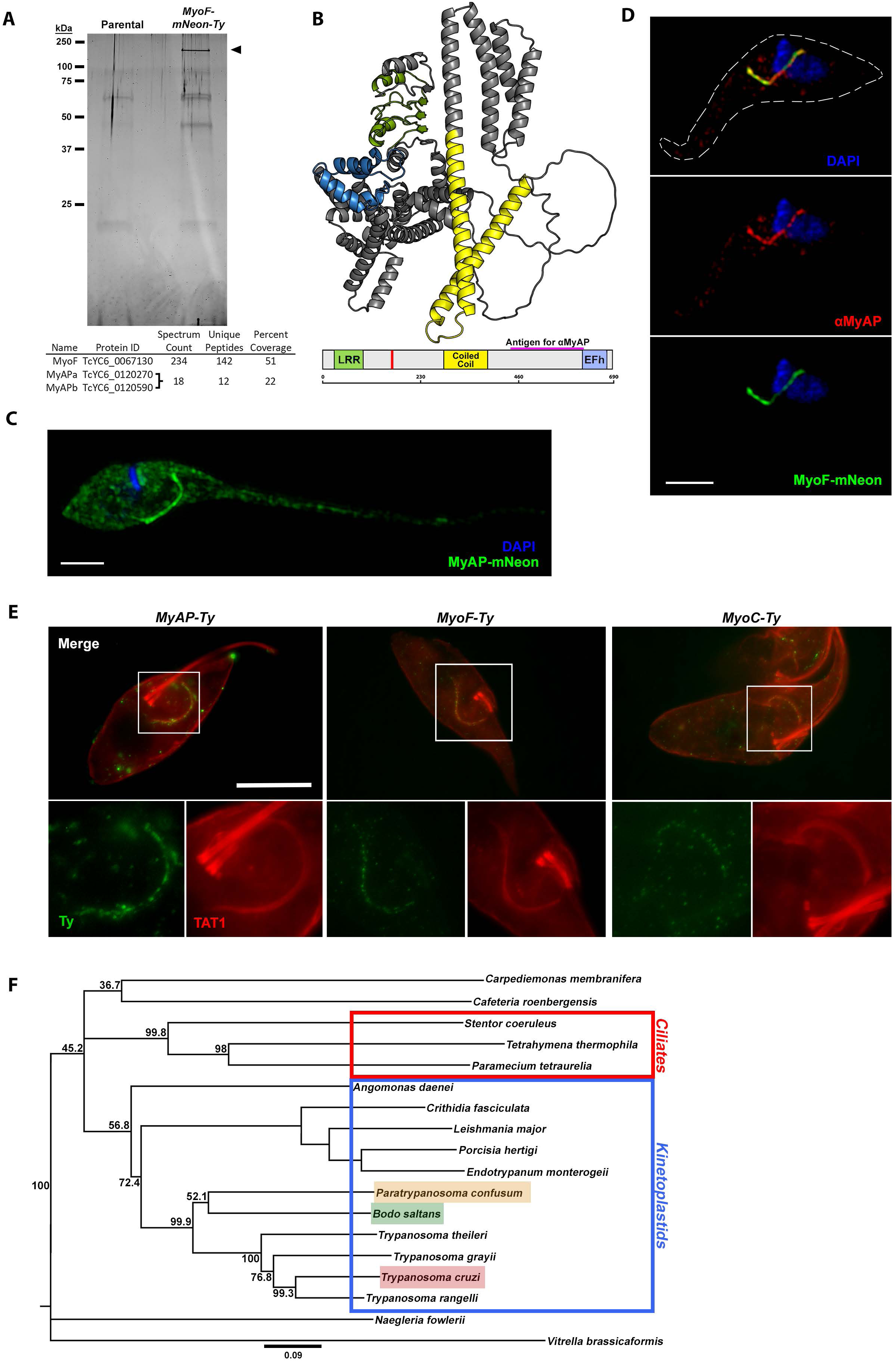
A Myosin Associated Protein (MyAP) Localizes to the SPC Microtubules. **A.** Oriole-stained gel (*top*) showing the eluate from αTy co-IP of MyoF-mNeon-Ty (*arrow*) overexpressing parasites. Table (*bottom*) shows mass spectrometry results from MyoF pulldown and the two identified isoforms of MyAP. **B.** MyAP AlphaFold predicted structure of (TcCLB.508479.180) (*top*) and predicted domains (*bottom*) on an amino acid scale line. Predicted leucine rich repeat (*green*), coiled coil (*yellow*), and EF-hand domains (*blue*) are shown. Also shown are the amino acids found only in the MyAP ***b*** isoform (*red vertical line*) and the antigenic region used for MyAP antibody generation (*pink horizontal line*). **C.** Fluorescence microscopy of a transiently overexpressed MyAP-mNeon showing localization to the typical SPC linear structure in epimastigotes. **D.** Immunofluorescence assay in MyoF-mNeon expressing epimastigotes showing co-localization between the SPC myosin MyoF-mNeon and αMyAP mouse antibody (1:200). **E.** Ultrastructure expansion microscopy reveals that MyAP (*top*), MyoF (*middle*), and MyoC (*bottom*) all localize to the SPC associated microtubules labelled by TAT1. **F.** Bootstrapped consensus tree shows the relationship between *T. cruzi* MyAP and its orthologs in a diverse range of protozoa including ciliates (*red box*) and other kinetoplastids (*blue box*). An ortholog from *Vitrella brassicaformis* was chosen as the outgroup. Scale bars 2 μm (C, D) and 20 μm (E).

To date, the vast majority of proteins we have identified as being targeted to the endocytic structure can be found only in SPC containing kinetoplastids. A broad phylogenetic analysis of the MyAP protein, however, has shown that orthologs of this protein can be found in a variety of protozoans including *Leishmania spp.* (**Figure 3F** phylogenetic tree and **Supplementary Figure S4** sequence alignment). Although *Leishmania spp.* lack an SPC and endocytose exclusively via the flagellar pocket, they nonetheless appear to retain a vestigial microtubule rootlet-like tract (similar to the CyQ/CyT) along which endocytosed material is trafficked to the cell posterior (30, 31) and it remains possible that the leishmanial MyAP ortholog may localize to this microtubule track as well. More broadly, however, the presence of MyAP orthologs in distantly related SPC containing species, including ciliates, highlights the potential for an evolutionarily conserved role in protozoan endocytosis.

### Loss of MyAP results in ablation of endocytosis and mistargeting of both MyoF and MyoC

To assess the functional role of MyAP in endocytosis, we implemented the CRISPR/Cas9 dependent gene deletion strategy, as performed previously with MyoF and MyoC. We deleted both paralogs of MyAP simultaneously using the same CRISRP targeting gRNA and drug selection cassette (**Figure 4A** methodology). We first generated a clonal cell line lacking both chromosomal copies of paralogs ***a*** and ***b*** of MyAP (*ΔMyAP*) and used this clone to, in turn, generate a complemented line with the subtype ***b*** paralog (*ΔMyAP::MyAP-Ty*). We verified disruption and restoration of the MyAP locus using diagnostic PCR (**Figure 4B**) and showed the loss and restoration of protein expression using our in-house derived MyAP antibody (**Figure 4C** Western blot). Following deletion of MyAP, we subjected parasites to our flow cytometry-based feeding assay and found that the deletion mutants demonstrated a complete lack of measurable endocytic activity while the complementation of MyAP-Ty fully restored endocytosis (**Figure 4D** and quantified in **4E**). The loss of MyAP and associated endocytic function also did not alter the growth of epimastigotes *in vitro* (**Supplementary Figure S1C and S1D**). In observing the essentiality of MyAP for endocytic function, we were curious if this impacted MyoF or MyoC overtly. We transfected the parental, null mutant (*ΔMyAP*) and complemented lines (*ΔMyAP::MyAP-Ty*) (**Figure 4F** left, middle and right panels respectively), with either the MyoF or MyoC-mNeon-Ty fusion constructs and examined myosin motor localization after 24 hours (hrs). Intriguingly, mutants lacking MyAP demonstrated an inability to properly target MyoF and MyoC to the canonical linear cytopharynx structure of the SPC and were instead found to be distributed diffusely throughout the cytosol (**Figure 4F** middle panels). MyAP complemented parasites restored normal localization of both motors (**Figure 4F** right panels). It is also worth noting that MyoB and MyoE were unaffected by the loss of MyAP and continued to be properly targeted to the pre-oral ridge region (**Supplementary Figure S5A**). Additionally, loss of MyoF or MyoC either separately or together did not impact the localization of MyAP suggesting that MyAP’s localization does not rely on the presence of myosin motors (**Supplementary Figure S6**). Using expansion microscopy, we were also able to show that loss of MyAP did not directly affect the cytostomal microtubules (CyQ: red arrow, and CyT: green arrow) (**Supplementary Figure S2F**). This data, therefore, suggests that MyAP plays a critical role in the specific recruitment of MyoF and MyoC to the SPC rootlet microtubules, thus potentially explaining why the severity of the observed endocytic defect mirrors the *ΔFΔC* knockout line.

**Figure 4.**
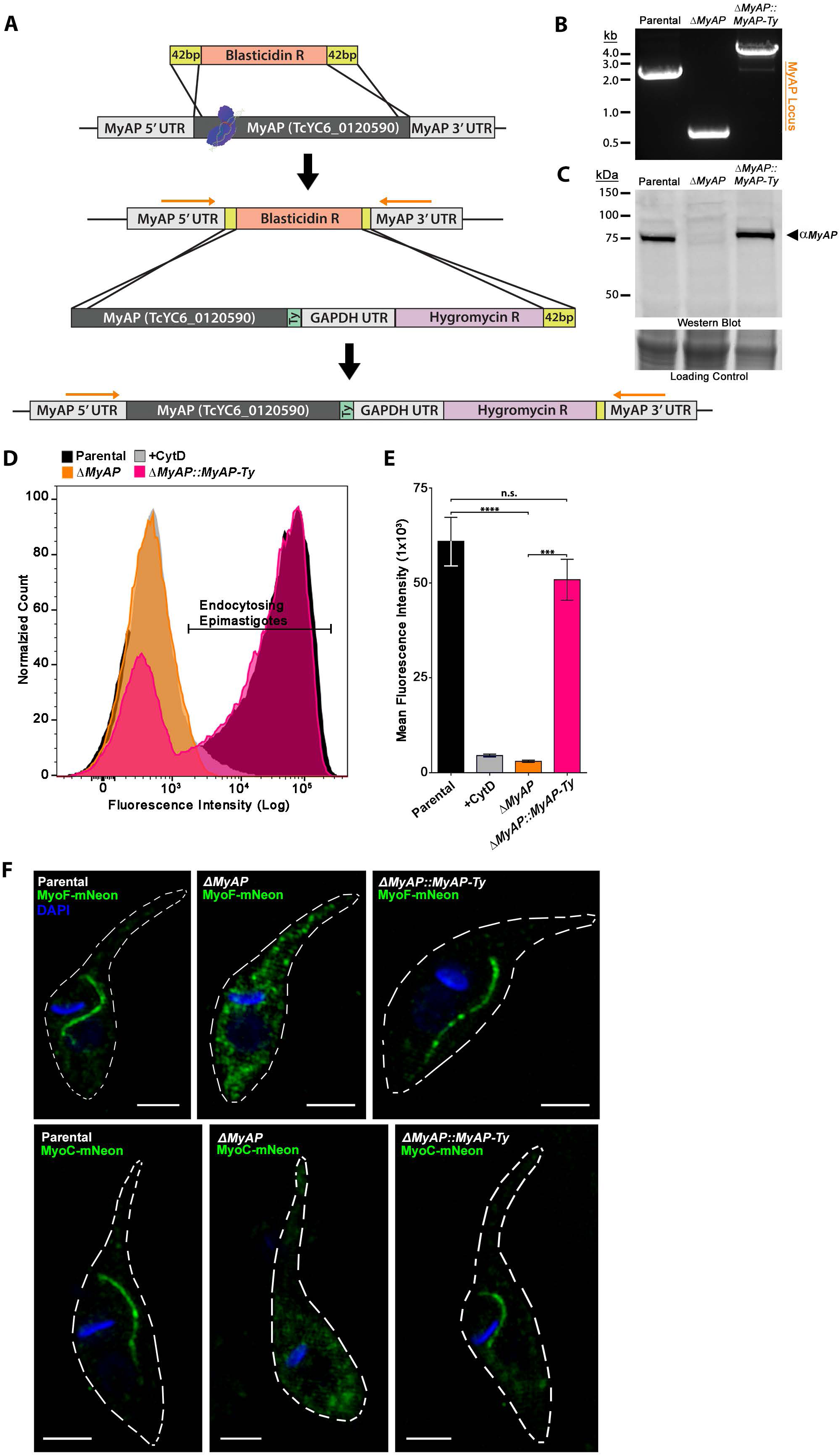
Deletion of MyAP Produces an Endocytic-Null Phenotype in Epimastigotes. **A**. Scheme for the CRISPR/Cas9 gene deletion and complementation of MyAP. Location of primers used for PCR verification are also shown (*arrows*). **B.** PCR amplification of the MyAP genomic locus shows replacement of both Parental loci (high-molecular-weight (MW) band) with the blasticidin resistance gene (low-MW band) in *ΔMyAP* mutants. Complementation with MyAP-Ty into the original MyAP locus (*ΔMyAP::MyAP-Ty*) is demonstrated by the high-MW band. **C.** Western blot of Parental, *ΔMyAP*, and *ΔMyAP::MyAP-Ty* epimastigote lysates using αMyAP mouse antibody (1:500) showing the successful deletion and restoration of MyAP protein. **D.** Flow cytometry analysis of fluorescent BSA fed epimastigotes shows an absence of detectable endocytosis in *ΔMyAP* epimastigotes (*orange*) similar to the Cytochalasin D treated negative control (*gray*). This endocytic defect is rescued in the *ΔMyAP::MyAP-Ty* complemented mutants (*magenta*). **E.** Quantification of three biological replicates of feeding assays, as shown in panel D, demonstrates ablation of endocytic activity in *ΔMyAP* epimastigotes and its rescue upon complementation (*ΔMyAP::MyAP-Ty*). **F.** Expression of MyoF-mNeon (*top*) and MyoC-mNeon (*bottom*) in Parental, *ΔMyAP*, and *ΔMyAP::MyAP-Ty* epimastigotes shows that MyoF and MyoC are mislocalized to the cytosol in the *ΔMyAP* mutants (*middle*). Localization to the SPC is rescued upon complementation of MyAP (*ΔMyAP::MyAP-Ty, right*). Scale bars 2 μm.

### The paired EF-hand structure is essential for MyAP function independent of its myosin motor recruitment activity

How the cytopharynx myosins (MyoF and MyoC) are targeted to the SPC and regulated remained an unknown aspect of this endocytic organelle’s function. With the discovery that MyAP is necessary for the proper localization of these motors, we aimed to dissect the functional contribution of the major identified LLR and EFh domains of MyAP to this activity (**Figure 5A** schematic). Using the endocytic null *ΔMyAP* line as the background strain, we first generated individual domain deletion constructs of MyAP lacking either the LRR or EFh domains and reintroduced this mutant gene into the endogenous locus in a manner analogous to that used in the original complementation line (see **Figure 4A** methodology). Clonal cell lines of both the *ΔMyAP::MyAPΔLRR-Ty* (green arrow) and *ΔMyAP::MyAPΔEFh-Ty* (blue arrow) complements were isolated and validated via diagnostic PCR (**Figure 5B**) and Western blot analysis of mutant MyAP protein expression (**Figure 5C**). We initially characterized the capacity of these complementation mutants to restore endocytic function and found that the loss of the LRR domain was not detrimental and endocytosis was restored to parental line levels (**Figure 5D** and quantified in **5E** (green)). In contrast, the EFh proved critical, as loss of this domain mirrored the phenotype of the parent *ΔMyAP* KO line (**Figure 5D** and quantified in **5E** (blue)). At this point, it was unclear how these deletion mutants were impacting myosin motor localization, so we again examined the targeting of MyoF-mNeon in both the LRR (*ΔMyAP::MyAPΔLRR-Ty*) and EFh deletion (*ΔMyAP::MyAPΔEFh-Ty*) lines. We found that neither a lack of an LRR nor EF-hand impacted proper targeting of MyoF-mNeon to the cytopharynx (**Figure 5F**). The EF-hand, therefore, appears dispensable for myosin recruitment, yet may still function to regulate the activity of the motor protein itself as is often the case with regulatory myosin light chain proteins (32). We next examined the functional contribution of three predicted calcium coordinating aspartic acid residues of the EF-hand through mutational analysis (D616A, D641A and D652A) (**Supplementary Figure S7A** EFh AlphaFold structure with aspartic acid residues highlighted). The resulting complementation mutants were validated via diagnostic PCR and Western blotting (**Supplementary Figure S7B** and **S7C**). Endocytic activity of the resulting complemented mutant lines was found to be indistinguishable from wild-type MyAP (**Supplementary Figure S7D**) (33–35). While the EFh domain in its entirety is essential for endocytic activity, our attempts to compromise this structure’s ability to bind calcium failed to disrupt endocytosis. This result suggests that either the EFh domain no longer binds calcium or alternatively that when calcium is bound it serves to negatively regulate motor function.

**Figure 5.**
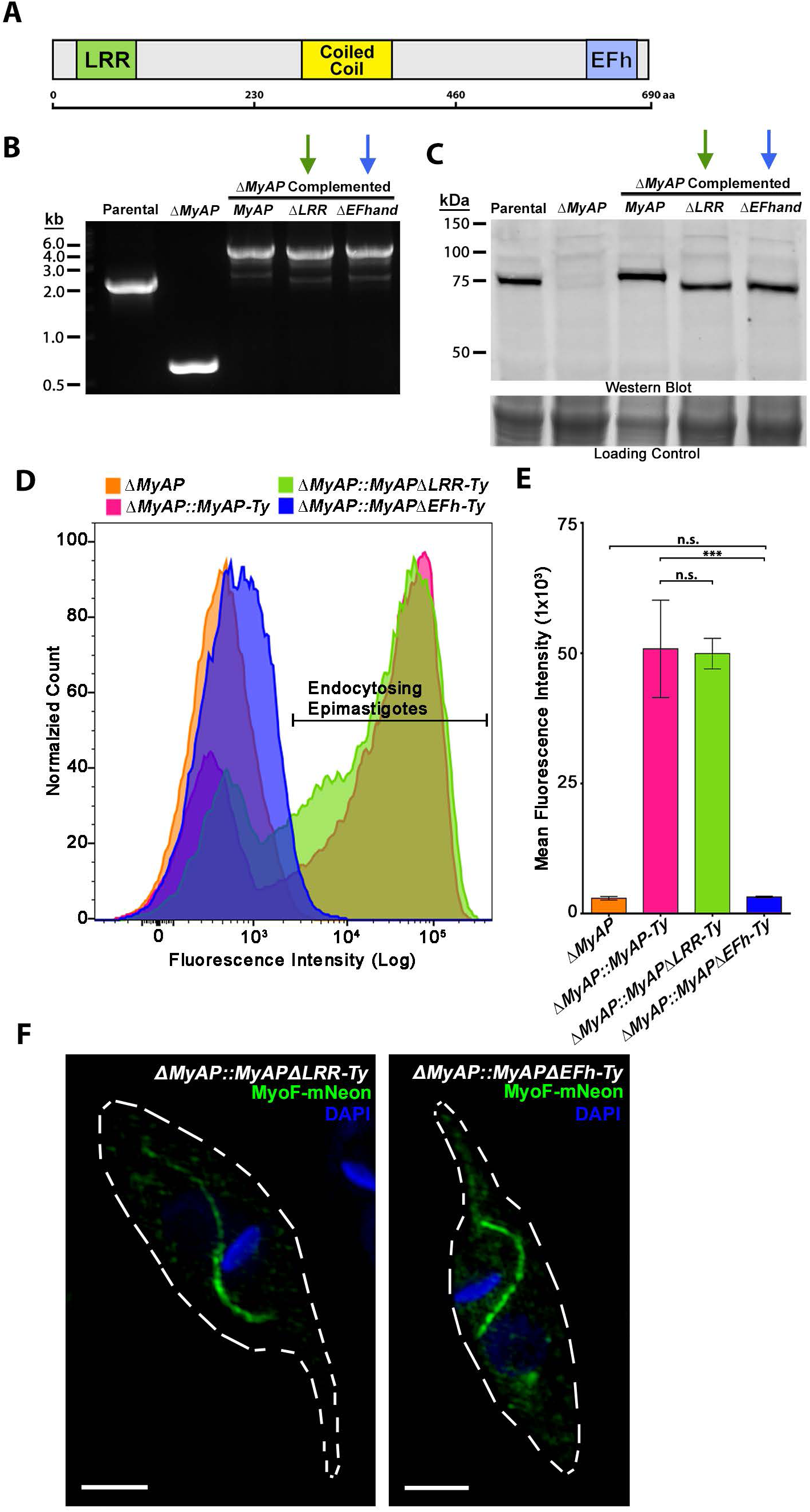
The Predicted Calcium Binding EF-hand Structure of MyAP is Essential for Endocytic Activity. **A.** Scheme of MyAP showing the predicted Leucine Rich Repeat (*green*) and paired EF-hand domains (*blue*) chosen for removal from the MyAP coding sequence prior to complementation into *ΔMyAP* epimastigotes. **B.** PCR amplification of the MyAP locus showing complementation of the various MyAP genes that were altered via mutagenesis. **C.** Western blot with αMyAP shows that the various MyAP versions are being expressed and translated in the complemented mutant epimastigotes. **D.** Flow cytometry analysis of fluorescent BSA fed epimastigotes shows a negligent restoration of endocytic activity in *ΔMyAP* epimastigotes complemented with MyAP lacking the EF-hand structure (*ΔMyAP::MyAP-EFh-Ty) (blue*). In contrast, normal endocytic activity was observed in epimastigotes complemented with MyAP lacking the LRR domain (*ΔMyAP:: MyAP-LRR-Ty*) (*green*). **E.** Quantification of three biological replicates of feeding assays, as shown in panel D, shows the failed restoration of endocytic rate in parasites complemented with *MyAP-EFh-Ty (blue*) when compared to those complemented with *MyAP-LRR (green*) or wild type *MyAP-Ty.* (*magenta*). **F.** Unlike what was observed in the *ΔMyAP* endocytic-null mutants, MyoF-mNeon localizes normally to the SPC in both *ΔMyAP::MyAP-LRR-Ty* and *ΔMyAP::MyAP-EFh-Ty* epimastigotes. Scale bars 2 μm.

### Loss of endocytosis reduces lipid uptake and storage in epimastigote reservosomes

Although the ablation of MyAP had a severe impact on endocytosis in *T. cruzi* epimastigotes, it was unclear as to how this was impacting the SPC or internal organellar structures, as no clear impact on growth *in vitro* had been observed (**Supplementary Figure S1C** and **S1D**). We subjected parental Y-strain and *ΔMyAP* KO parasites first to scanning electron microscopy (SEM) in order to assess changes to the cytostome opening. We found that the loss of MyAP had no overt effect on the overall size and appearance of the cytostome at the surface of these parasites (**Figure 6A**). We next examined the internal structures of the parasite and employed transmission electron microscopy (TEM) to compare parental, *ΔMyAP* KO and *ΔMyAP::MyAP-Ty* complemented epimastigotes and observed that endocytic-null mutants demonstrated a marked difference in the composition of their pre-lysosomal reservosome storage structures (**Figure 6B** middle: orange panel green outlined vesicles). Although still present, the reservosomes of the *ΔMyAP* mutant lacked the typical electron lucent voids indicative of lipid inclusions and thus appeared uniformly electron dense (36). This suggested that parasites were no longer able to accumulate lipids in their reservosomes via endocytosis. Complementation of MyAP in the deletion strain (*ΔMyAP::MyAP-Ty*) restored both endocytic capacity and the presence of the lipid inclusions (**Figure 6B** right: red outlined panel and green outlined vesicles). To obtain a direct look at the parasite lipidome, we analyzed the compositional changes in sterol makeup of endocytic null mutants. We subjected Parental, *ΔMyAP* and *ΔMyAP::MyAP-Ty* complemented epimastigote parasites, grown to log phase *in vitro*, to MS based lipidomics analysis. The sterol composition of fresh epimastigote LIT/LDNT media was also assessed in parallel (37). A principal component analysis of the dataset demonstrated the significant differences in the *ΔMyAP* lipid profiles as compared to the parental and *ΔMyAP::MyAP-Ty* complemented lines (**Supplementary Figure S8A**). We first compared the main, endogenously synthesized, parasite specific sterol (ergosterol) to the exclusively scavenged host-derived, or in this case media derived, sterol (cholesterol) (20, 38). In this analysis we found that while endogenously synthesized ergosterol levels were unchanged in all three parasite strains regardless of endocytic capacity, scavenged cholesterol was dramatically reduced only in the endocytic-null mutant, with levels being restored in the complemented line (**Figure 6C**). This analysis further supports the myriad observations we have presented here, demonstrating that these endocytic-null mutant parasites lack the ability to actively phagocytose via their SPC and thus fail to accumulate exogenous nutrients into their reservosomes.

**Figure 6.**
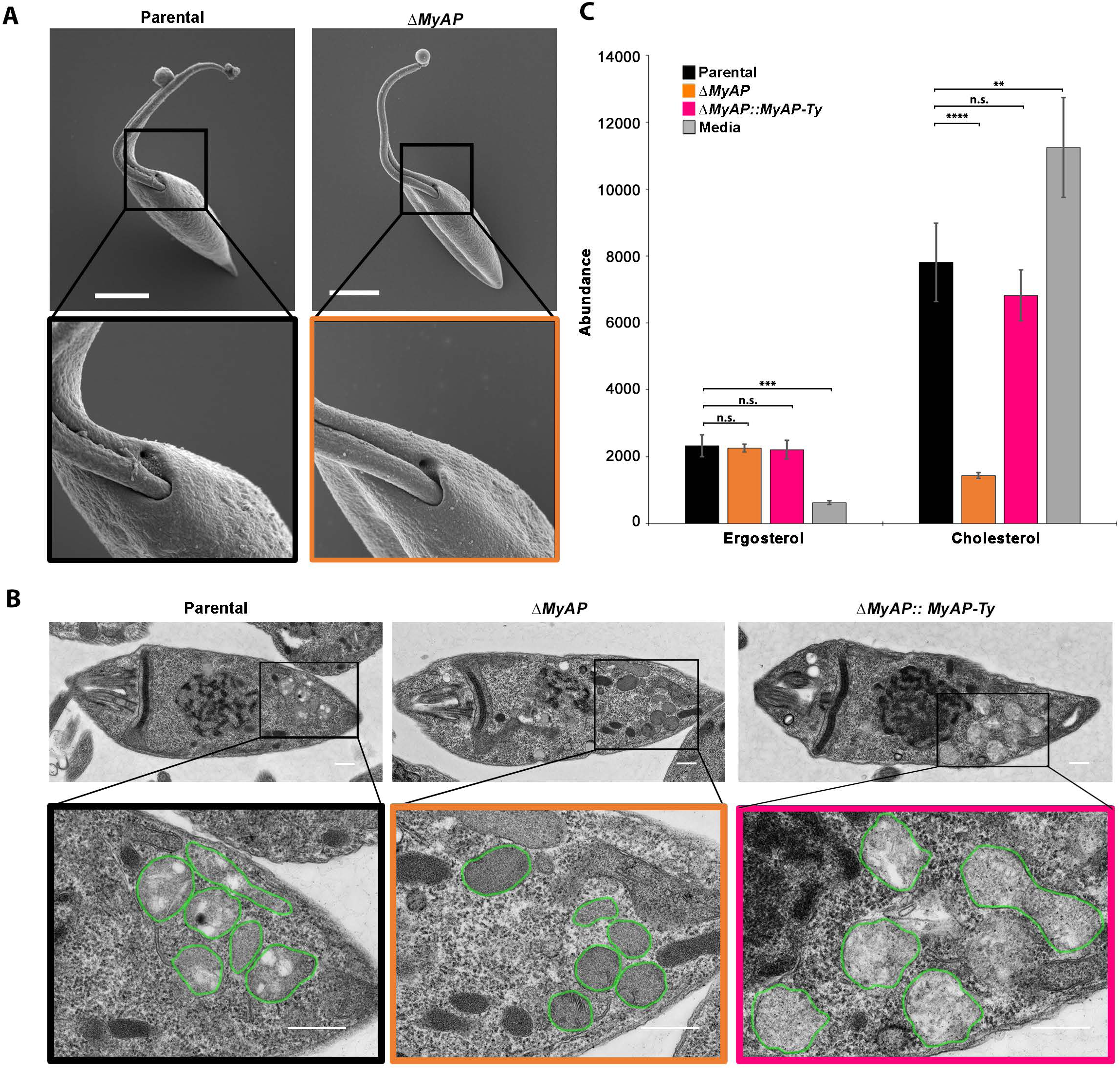
Deletion of MyAP Results in Altered Reservosomes and Defects in Sterol Accumulation. **A.** Scanning electron micrographs reveal that the endocytic-null phenotype of *ΔMyAP* epimastigotes is not caused by the lack of a cytostome opening, as it can be observed in both the Parental and *ΔMyAP* epimastigotes. Scale bars 2 μm. **B.** Transmission electron microscopy of Parental Y Strain epimastigotes (*left*) shows electron lucent structures in the pre-lysosomal reservosomes that have the typical appearance of typical of lipid inclusions. These inclusions are absent in the uniform electron dense reservosomes of *ΔMyAP* epimastigotes (*middle*) and their presence is rescued upon complementation (*right*). Scale bars 0.5 μm. **C.** Lipidomic analysis of the sterol content in *ΔMyAP* epimastigotes (*orange*) shows a dramatically reduced cholesterol content (*right*) compared to the Parental (*black*) that is restored upon complementation (*magenta*). No such difference is observed in the levels of endogenously synthesized ergosterol (*left*).

## Discussion

In our previously published study, we identified four distinct orphan myosin motors that are targeted to the oral apparatus of *Trypanosoma cruzi* and characterized the contribution of the cytopharynx targeted MyoF to SPC mediated endocytosis (19). Here we have continued this dissection of the molecular machinery responsible for *T. cruzi* endocytosis by focusing on the two cytopharynx localized myosins: MyoF and MyoC. While the deletion of MyoF alone significantly reduced the rate of endocytosis, the combined deletion of both motors (*ΔFΔC*) produced endocytic-null parasites that lacked detectable levels of protein uptake. The ablation of endocytosis, again, had no significant effect on the growth rate of *T. cruzi*, further supporting the notion that this process is not essential for viability under laboratory culture conditions (19).

To begin expanding our understanding of the protein complexes contributing to SPC function, we carried out a co-IP of overexpressed MyoF-mNeon-Ty in insect stage epimastigotes followed by mass spectrometry analysis. From this, we identified a protein containing both an N-terminal leucine rich repeat (LRR) and a putative C-terminal calcium responsive paired EF-hand (EFh) structure which we refer to as the <u>m</u>yosin <u>a</u>ssociated <u>p</u>rotein or MyAP. This protein was chosen for further study due in part to the presence of the paired EF-hand domain which we suspected might interact with the IQ motifs found on MyoF to possibly regulate motor function (39, 40). Upon closer examination of the Y-strain genome (24), we also discovered that MyAP existed as two distinct paralogs (which we refer to as subtype ***a*** and ***b***) distinguished solely by the presence of a six amino acid insertion. We first confirmed MyAP’s SPC localization using IFAs by either overexpressing it as an mNeon fusion protein or staining the endogenous protein with our in-house generated MyAP antibody. Both techniques demonstrated a clear localization of MyAP to the SPC cytopharynx structure. However, because the cytopharynx complex itself is composed of both a dynamic membrane tubule as well as adjacent stable microtubule rootles, we set out to determine with greater precision the location of MyAP within this region of the endocytic apparatus itself. With the aid of expansion-based immunofluorescence microscopy, we were able to show that MyAP is fixed upon the microtubule rootlet structure, rather than on the more dynamic membrane tubule. This technique also allowed us to examine the localization of MyoF and MyoC and we found both motors also localized on these root fibers beginning at the cytostome entrance. The targeting of MyoF to these microtubules validates prior observations by Alves et al. who, by using immuno-EM of epitope tagged MyoF on membrane extracted *T. cruzi* cytoskeletons, demonstrated that MyoF associates with SPC rootlet microtubules ((28) and reviewed in (12)). Our subsequent direct deletion of both MyAP paralogs using CRISPR/Cas9, resulted in endocytic-null parasites that phenocopied our myosin double deletion mutants (*ΔFΔC*) and we were able to completely restore this endocytic defect via complementation. The essentiality of MyAP to endocytosis and its presence in a number of distantly related SPC containing protozoans leads us to believe that this protein may play an ancestrally conserved role in SPC mediated protozoal phagotrophy. To address the means by which MyAP was impacting endocytosis, we first localized MyoF and MyoC in the *ΔMyAP* line and, intriguingly, we found that the cytopharynx motors were cytosolic and no longer associated with the SPC. Proper targeting of MyoF/C was again restored when MyAP was complemented back into the KO. It, therefore, appeared likely that the reason the loss of MyAP phenocopies the *ΔFΔC* endocytic defect is due, in part, to the inability to recruit MyoF/C to the cytostomal microtubules, a seemingly necessary step in facilitating endocytosis. What this work has not yet demonstrated, however, is if this interaction between MyAP and the motors is direct or if there are additional protein factors involved which mediate these interactions. With respect to the overall model of SPC function, these observations suggest that the movement of endocytosed membrane into the cell cytosol is propelled by a linear array of myosin motors fixed onto rootlet microtubule structures. We reason that since these motors must walk on actin filaments, likely in a plus-end directed fashion, the actin polymers themselves must be associating with the cytopharynx membrane in a polarized orientation. This model, therefore, predicts that the polymerization of actin is likely initiated at the cytostome entrance in order to coat the emerging cytopharynx membrane with polarized microfilaments that the fixed myosins can pull on unidirectionally (reviewed in (12)).

In examining the functional role of the two identified domains found in MyAP (LRR and EFh) we reintroduced, via complementation, the MyAP gene lacking either the LRR domain or the EFh. While the LRR domain appeared fully dispensable for normal protein function, loss of the EFh failed to restore endocytosis. We initially hypothesized that the EFh was playing a role in myosin motor recruitment but, to our surprise, we found that the *MyAPΔEFh-Ty* mutant restored proper MyoF targeting. This implied to us that the MyAP protein is potentially performing two independent yet necessary functions: recruitment of myosin motors to the SPC microtubules on the one hand and regulation of myosin motor activity via the EFh on the other. To examine this possibility in more detail, we next mutated three predicted calcium coordinating aspartic acid residues of the EFh and found that none of these mutations negatively impacted endocytosis. Although we have yet to directly confirm the calcium binding capability of this EFh biochemically, our mutations of conserved residues indicate that the structural fold of the calcium-free EF-hand is sufficient to activate motor activity. This data also implies that if calcium binds to this domain, it likely promotes EFh disengagement leading to shutdown of the motor in a manner analogous to other EF-hand containing calmodulin myosin light chains (41–43). If this is the case, our results simply mirror prior studies where calcium binding mutants of calmodulin are unable to dissociate and lead to myosin motors which are constitutively active. A more complete understanding of the role that the MyAP EFh has on regulating MyoF/C motor activity will, in all likelihood, require biochemical analyses of purified proteins to characterize the nature of binding and activation of myosin activity by MyAP. More broadly, this observation raises the interesting prospect that calcium signaling itself may play an important role in regulating *T. cruzi* endocytosis.

In the absence of a clear *in vitro* growth defect exhibited by our endocytic null mutants, we continue to be faced with the central question regarding the ultimate role of endocytosis in this parasite’s life cycle. Although we have speculated extensively in our prior reports as to why the SPC was retained specifically in those trypanosomatids that are transmitted via the insect feces, we still lack definitive evidence showing when or where endocytosis is essential in the parasite life cycle (12, 19). However, what we have been able to demonstrate here using TEM and MS-based lipidomics analyses, is that these mutants have a dramatically reduced capacity to take in and store extracellular material such as host-derived cholesterol in their reservosome structures. Prior reports have shown that reservosomes play an important role in nutrient provisioning under starvation conditions since they transition from storage depots into digestive lysosomes to fuel the metabolic needs of metacyclogenesis, a necessary step in transmission (44–46). It remains possible that, when lacking endocytosis, parasites will present markedly reduced fitness under specific nutrient stress conditions. Future detailed parasite-vector studies will be needed to examine the ability of these endocytic-null parasites to effectively colonize the insect intestine, compete for nutrients and space with the associated insect microbiome and ultimately undergo metacyclogenesis as they prepare to infect their eventual vertebrate host. These studies will be the necessary next step if we are to provide a definitive answer as to why *T. cruzi* retained this ancestral feeding structure.

In conclusion, it is worth restating that the endocytic organelle which we have discussed and referred to here as the cyto<u>s</u>tome-cyto<u>p</u>harynx <u>c</u>omplex or SPC, is a fascinatingly complex and ubiquitous feeding structure found in vast numbers of heterotrophic protozoans which populate the world’s various ecosystems. In spite of both its presence in myriad protozoan species that play critical roles in environmental nutrient cycling and human disease, we know next to nothing about how the SPC is formed or functions at a mechanistic level. Using the human kinetoplastid parasite *Trypanosoma cruzi* as a model, our group has been able to initiate a systematic dissection of the structural and mechanistic underpinnings of the SPC organelle. By taking advantage of the fact that *T. cruzi* is both genetically tractable and does not require SPC function for *in vitro* growth we can examine the molecular components of this organelle without negatively impacting parasite fitness in culture. This, in the end, is a fortuitous finding since *T. cruzi* continues to lack the necessary genetic tools to study the function of essential genes (47). An array of important questions regarding the formation and regulation of the SPC organelle clearly still remain and future studies will no doubt continue to provide important insight into the molecular mechanisms undergirding this fundamental process of protozoan phagotrophy.

## Acknowledgments

We would like to sincerely thank the members of the Center for Tropical and Emerging Global Diseases (CTEGD), Julie Nelson and the CTEGD Cytometry Shared Resource Laboratory, the T32 Training in Tropical and Emerging Global Diseases grant (T32AI060546) and funding from the NIH (R01AI163140 and R01GM144545).

## Materials and Methods

### Parasite cultures

Y-strain epimastigotes were cultured in LIT/LDNT medium (37) supplemented with 15% 76°C heat-inactivated fetal bovine serum (FBS) (VWR, USDA certified). To obtain amastigotes, metacyclic trypomastigotes generated via epimastigote starvation as previously described (18) were added to T25 flasks or coverslips containing confluent human foreskin fibroblasts (HFF). Infected monolayers were maintained in high-glucose Dulbecco’s modified Eagle’s medium (DMEM-HG) (HyClone) supplemented with L-glutamine and 1% 56°C heat-inactivated Cosmic Calf serum (CCS) (HyClone).

### Epimastigote growth assays

Standard growth assays were performed as previously described (18). 5.0 x 10^6^/ml epimastigotes of each line were seeded into 2 mL LIT media containing 15% of 76°C heat-inactivated FBS in a T12.5 flask. Counting every 24 hrs was performed using a Z1 Coulter counter (Beckman Coulter) counting at a 1:200 dilution and averaging three counts per sample. Cumulative growth assays were performed as described above except that the parasite concentration was reset to 5.0 x 10^6^/ml each day with fresh media after counting.

### Transfections and overexpression

Transfections were performed as previously described (48) with some modifications. Precipitated DNA pellets were initially resuspended in 10 μL nutrient free TAU containing 5% 76°C heat inactivated FBS, prior to adding cells with in electroporation buffer. 2.5 x 10^7^ epimastigotes were transfected using two pulses of the BTX ECM 830 system (Harvard Apparatus) for all transfections. A list of primers used during this work can be found in Table S1 in the supplemental material. For overexpression of MyoC-mNeonTy and MyoF-mNeonTy in MyAP knockout epimastigotes, the plasmids generated in our previous work (19) were used.

### Co-immunoprecipitation and liquid chromatography/mass spectroscopy

Co-immunoprecipitation of overexpressed MyoF-mNeonTy using an anti-Ty antibody was performed as previously described (18, 49). Eluates were run briefly on an Any-kD SDS-PAGE gel (BioRad) to separate out most of the Ty peptide used for the elution process and then the lane was excised prior to washing in 50% methanol for LC/MS analysis. The samples were analyzed via data-dependent electrospray LC-MS/MS on a Thermo Q-Exactive Orbitrap mass spectrometer. Trypsin was selected as the protease, with maximum missing cleavage set to 2. A 1% false discovery rate cutoff was selected for peptide, protein, and site identifications. MS results were searched against the Y-strain YC6 genome (tritrypdb.org).

### Gene deletion and complementation using CRISPR/Cas9

Deletion and complementation were performed as previously described (19, 48) with modifications. G418 (1,000 μg/ml) added 24 hrs after Cas9 transfections was only maintained for 3 days when using Blasticidin repair templates for gene deletions and 5 days when using Hygromycin repair templates for complementation, to help stall the growth of untransfected epimastigotes immediately after transfection. Other aspects of this methodology including plasmids used (pTMiniTrex), gRNA design, and subcloning were consistent with the previously published method cited above. Double gene deletion was performed sequentially using a Blasticidin resistance repair template for MyoF and a Hygromycin resistance repair template for MyoC. Complementation of double knockouts utilized the G418 resistance cassette and selection with only 250 μg/mL G418 after the initial 72 hrs (during which the normal 1000 μg/mL was used). Mutagenesis of complement templates was performed using the Q5 Mutagenesis kit (NEB) with primers referenced in the primer table S1.

### Western blotting and fluorescent microscopy

Western blotting was performed as previously described(18). Mouse anti-MyAP antibody was used at 1:500. Fluorescence microscopy was performed as previously described with modifications for the MyAP antibody images (18). A modified fixation protocol was used for the MyAP antibody images to improve MyAP labeling. 1×10^7^ epimastigotes were pelleted at 1,000 x g for in a 1.5 mL tube for 3 min. LIT/LDNT culture media supernatant was removed and then, without disturbing the pellet, the epimastigotes were fixed with a rapid addition of 1 mL of −20°C Methanol and disruption of the pellet. Fixation was continued at −20°C for 5 min then transferred to −80°C for an additional 10 min to maintain cold temperature during the proceeding spin. Fixed parasites were spun down in a 4°C centrifuge at 1,500 x g for 3 min, then washed three times in 1 mL pH 7.4 PBS using the same 1,500 x g spins. ConA labeling and the remaining IFA steps were performed as previously described (18). Mouse polyclonal anti-MyAP was used in IFAs at a 1:200 dilution. Rabbit anti-cruzipain antibody (used at 1:1000) was a generous gift from Roberto Docampo.

### Flow cytometry-based endocytosis assays

Endocytosis assays were performed as previously described (19). 5 x 10^6^ log phase epimastigotes of each line were pelleted in 1.7 mL centrifuge tubes, washed in 500 μ? HBSS (Hanks balanced salt solution) and either treated with 10 μM Cytochalasin D or mock treated for 10 min. Epimastigotes were then incubated with Alexa Fluor 647-conjugated BSA for 30 min at 28°C, washed in 10 mL HBSS then resuspended in 1 mL HBSS before being run on a Quanteon flow cytometer (Acea Bio). Analysis was performed using FlowJo software.

### Sequence alignment, tree generation and structure prediction

Alignments were performed using the T-Coffee alignment server (50) and alignment figures were generated using Jalview 2 software (51). Consensus tree of predicted MyAP orthologues (identified via NCBI Blast (52)) was generated in the Geneious Prime software suite using the Jukes-Cantor Genetic Distance Model, Neighbor-Joining tree build method, and Bootstrapped. Structure prediction of MyAP was obtained from the AlphaFold database (https://alphafold.ebi.ac.uk/entry/Q4E2K6) (53) and figures generated using Pymol software (Schrodinger).

### Transmission electron microscopy

For morphological analyses at the ultrastructural level, parasites were fixed in 2% paraformaldehyde/2.5% glutaraldehyde (Ted Pella Inc., Redding, CA or other source) in 100 mM sodium cacodylate buffer, pH 7.2 for 2 hr at room temperature. Samples were washed in sodium cacodylate buffer at room temperature and post-fixed in 2% osmium tetroxide (Ted Pella Inc) for 1 hr at room temperature. Samples were then rinsed in dH20, dehydrated in a graded series of ethanol, and embedded in Eponate 12 resin (Ted Pella Inc). Sections of 95 nm were cut with a Leica Ultracut UCT ultramicrotome (Leica Microsystems Inc., Bannockburn, IL), stained with uranyl acetate and lead citrate, and viewed on a JEOL 1200 EX transmission electron microscope (JEOL USA Inc., Peabody, MA) equipped with an AMT 8 megapixel digital camera and AMT Image Capture Engine V602 software (Advanced Microscopy Techniques, Woburn, MA).

### Scanning electron microscopy

Epimastigotes were fixed in a freshly prepared solution of LIT/LDNT media with 2% paraformaldehyde/2.5% glutaraldehyde for 1 hr at room temperature. Post-fixation, SEM samples were gently pelleted and washed in 0.15 M cacodylate buffer. This was repeated two additional times after which the samples were incubated with 1% osmium tetroxide in 0.15 M cacodylate buffer for 45 min in the dark. The samples were then gently pelleted and washed in ultrapure water. After three rinses, pelleted samples were resuspended in 100 μl of ultrapure water and loaded onto coverslips freshly coated with 1 mg/ml poly-L-lysine. Cells were allowed to settle and attach to the coverslips for an hour. Samples were then dehydrated in a graded ethanol series (10%, 30%, 50%, 70%, 90%, 100% x 3) for 10 min each step. Following dehydration, the samples were loaded into a critical point drier (Leica EM CPD 300, Vienna, Austria) which was set to perform 12 x CO2 exchanges at the slowest speed. Once dried, coverslips were mounted on aluminum stubs with carbon adhesive tabs and coated with 10 nm of carbon and 10 nm of iridium (Leica ACE 600, Vienna, Austria). SEM images were acquired on a FE-SEM (Zeiss Merlin, Oberkochen, Germany) at 1.5 kV and 0.1 nA.

### Ultrastructure expansion microscopy

For each expansion gel, a volume of 2x fixative was added to 1 x 10^6^ cells in an equal volume of media to provide a 1x final concentration of fixatives: 0.7% paraformaldehyde (Thermo Fisher Scientific), 1% acrylamide (Bio-Rad, Hercules, CA), in 1x PBS (Fisher Scientific, Pittsburgh PA). Cells were harvested by centrifugation at 1000 x g for 10 min, then washed with 1x fixative (0.7% paraformaldehyde, 1% acrylamide, 1× PBS) and the centrifugation step repeated. Cell pellets were resuspended in 1x fixative and were briefly spun onto coverslips at 800 × g for 5 sec. The slow speed and short time preserve the morphology of the cells. The coverslips were inverted onto 80 μL droplets of 1x fixative in a humidified chamber for 3.5 hr at 37°C. Coverslips were then inverted into gelation solution (19% sodium acrylate (Pfaltz & Bauer, Waterbury CT), 10% acrylamide, 0.1 % Bis (Bio-Rad), in 1x PBS) activated with 0.5% TEMED (Bio-Rad) and 0.5% ammonium persulfate (Sigma-Aldrich, St. Louis, MO). Gelation reaction was allowed to solidify for 1 hr at 37°C. Gels on coverslips were incubated in denaturation buffer (200 mM sodium dodecyl sulfate (Fisher Scientific), 200 mM sodium chloride (Fisher Scientific), 50 mM TRIS Base (Fisher Scientific)) for 15 min at RT. Gels were then carefully peeled away from coverslip into a 1.6 mL microcentrifuge tube and then incubated in denaturation buffer at 95°C for 30 min. Gels were incubated in large petri dishes filled with deionized water to expand. A water change was done after 30 min and the gels were then allowed to expand fully overnight. The following day, the gels were shrunk in 1x PBS for 30 min. Shrunken gels were incubated in primary antibodies in 2% BSA (Fisher Scientific) in 1x PBS for 4 hr. Gels were washed 3 times in 1x PBS with 0.1% Tween-20, for 20 min each wash. Gels were then incubated in secondary antibodies in PBS with 2% BSA for 4 hr followed by three 20 min washes in 1x PBS with 0.1% Tween-20. Gels were transferred to large petri dishes filled with deionized water for expansion. After 30 min, the water was exchanged with fresh deionized water and the gels were allowed to fully expand overnight. The next day, gel punches were imaged on glass bottom microwell dishes (MatTek Corporation, Ashland, MA). Expansion microscopy images were taken on a Zeiss Axio Observer.Z1 microscope (Carl Zeiss Microscopy, Oberkochen, Germany) using a 100x/1.4 NA Plan Apochromat oil objective lens imaged with an ORCA-Flash 4.0 V2 CMOS camera (Hamamatsu—Shizuoka, Japan). Images were acquired in Slidebook 6 software (3i, Denver, CO) as z-stacks with a step size of 250 nm. Microscopy images were visualized with ImageJ (National Institutes of Health, Bethesda, MD), and figures prepared in Adobe Photoshop and Illustrator (CC 2022). Antibodies for expansion microscopy were sourced and diluted as follows: anti-Ty1 antibody (1:10) from Cynthia He (National University of Singapore, Singapore), anti-TAT1 antibody (1:100) from Jack Sunter (Oxford Brookes University—Oxford, United Kingdom). Secondary antibodies Goat anti-mouse IgG1 Alexa Fluor 488 (Thermo Fisher Scientific, Waltham MA) and goat anti-mouse IgG2a Alexa Fluor 568 (Thermo Fisher Scientific) were used at 1:100 dilution.

### Detergent fractionation for evaluation of cytoskeleton association

Detergent fractionation was performed similar to described by Alves et al. (28) with modifications. 30 x 10^6^ epimastigotes were washed 1x with cold Brinkley Buffer 1980 (BRB80, 80mM PIPES pH 6.8, 1 mM MgCl2, 1 mM EGTA, 1 mM EDTA) + 0.1 mM EDTA and Complete Protease Inhibitors (Roche). Detergent treatment was performed by adding 1% NP40 in BRB80 for 10 min at 4°C. After Centrifugation at maximum speed, the supernatant was spun down a second time and removed to a separate tube as the detergent soluble fraction. The detergent insoluble pellet was washed twice with 1 mL of BRB80 buffer +1% NP40 before being resuspended and boiled in SDS PAGE Buffer for SDS PAGE and immunoblot as described above.

### Statistical analysis

Statistical analysis was performed using the Prism Software suite unpaired *t*-test function. Three biological replicates were performed for each analyzed experiment. *P* values are denoted as follows: *, *P*<0.05; **, *P*<0.01; ***, *P*< 0.001; ****, *P* < 0.0001.

### Lipidomic analyses

Lipids were extracted from *T. cruzi* samples using a modified Bligh & Dyer method (54, 55). Dried lipid extracts were reconstituted in 500 μL of 1:1 chloroform/methanol. For lipidomic analysis, a 10x dilution of the lipid extract was prepared in 95% acetonitrile/5% water with 5 mM ammonium acetate. Lipids were separated on a hydrophilic interaction liquid chromatography column (Waters CORTECS HILIC, 2.1 x 100 mm, 1.6 μm) using a 12 min gradient of 95% acetonitrile/5% water with 5 mM ammonium acetate and 50% acetonitrile/50% water with 5 mM ammonium acetate (56). IM-MS measurements were collected using a Waters Synapt XS in positive and negative ionization modes with data-independent MS/MS collection. Lock-mass correction, peak picking, alignment, and normalization was performed with Progenesis QI (Nonlinear Dynamics) and multivariate statistical analysis was performed in EZinfo (Umetrics). Lipid identifications were assigned first to the class level based on annotated HILIC retention times of yeast total lipid extract, followed by species level annotations against an in-house version of LipidPioneer using a mass accuracy threshold of 10 ppm (57). A 15 min isocratic reversed-phase liquid chromatography (Waters CORTECS C18, 2.1 x 100 mm, 1.7 μm) method was used to confirm the identifications of cholesterol and ergosterol against reference standards (58). HPLC grade solvents (water, acetonitrile, methanol, and chloroform) and ammonium acetate were purchased from Thermo Fisher Scientific. Lipid reference standards were purchased from Avanti Lipids (Yeast Total Lipid Extract and d7-cholesterol) or Cayman Chemical (ergosterol).

## Author Contributions

RDE, NMC, MGE, PC, PG, KB and KH designed and performed the experiments, analyzed the data and generated the figures. RDE and NMC wrote the manuscript with co-author input.

## Declaration of Interests

The authors declare no conflict of interest.

---

**Supplementary Figure 1 Endocytic-Null Mutants Do Not Show a Significant Growth Defect in Culture**

**A, B.** Similar growth rate is seen in both cumulative (A) and absolute (B) growth rate assays of MyoF and MyoC knockout and complement mutant epimastigotes.

**C, D.** Likewise, no significant growth rate reduction can be seen in the endocytic-null *ΔMyAP* or *ΔMyAP::MyAP-EFh* epimastigotes in either cumulative (C) or absolute (D) growth rate assays.

**Supplementary Figure 2 Deletion of MyAP Does Not Alter the Localization of the Preoral Ridge Myosins**

Preoral ridge myosins MyoB (*left*) and MyoE (*right*) maintain their normal localization the preoral ridge in *ΔMyAP* epimastigotes.

**Supplementary Figure 3 Additional Expansion Microscopy Images Showing Microtubule Localization of MyoF, MyoC, and MyAP**

**A, B, C.** Additional supporting expansion microscopy showing TAT1 (*red*) labelled microtubule localization of MyAP-Ty, MyoF-Ty, and MyoC-Ty as in Figure 3 Panel E.

**D.** Negative control showing background Ty labeling in Parental line.

**E, F.** Expansion ultrastructure microscopy images showing TAT1 labeling of microtubules in *ΔMyAP* epimastigotes. In panel F, both the microtubule quartet (*red arrowhead)*) and triplet (*green arrowhead*) can be clearly differentiated, suggesting there is no disruption in microtubule formation.

**Supplementary Figure 4 MyAP is Enriched in the Detergent Insoluble Cytoskeletal Fraction**

Immunoblot of detergent extracted lysates show an enrichment of MyAP in the detergent resistant cytoskeletal fraction of the Parental and *ΔMyAP::MyAP-Ty* lines (*top*). Cruzipain was used as a marker for the detergent soluble fraction (S). The total absence of cruzipain labeling in the detergent insoluble cytoskeletal fraction (P) is evidence of a clean fraction (*middle*). Total protein stain from the BioRad Stain-Free gel used for this blot is also shown (*bottom*).

**Supplementary Figure 5 T-coffee Multiple Sequence Alignment and Gene IDs Used for Tree in Figure 3**

**Supplementary Figure 6 Localization of MyAP in Strains Used in This Study**

**A.** Immunofluorescence assay of MyAP using the αMyAP mouse antibody (1:200) in the Parental strain (green). ConA lectin surface staining of epimastigotes in red. DAPI in blue.

**B.** Immunofluorescence assay of MyAP using the αMyAP mouse antibody (1:200) in the *ΔMyAP* strain (green). ConA lectin surface staining of epimastigotes in red. DAPI in blue.

**C.** Immunofluorescence assay of MyAP using the αMyAP mouse antibody (1:200) in the *ΔMyAP*::*MyAP-Ty* strain (green). ConA lectin surface staining of epimastigotes in red. DAPI in blue.

**D.** Immunofluorescence assay of MyAP using the αMyAP mouse antibody (1:200) in the *ΔMyAP:: MyAPΔLRR-Ty* strain (green). ConA lectin surface staining of epimastigotes in red. DAPI in blue.

**E.** Immunofluorescence assay of MyAP using the αMyAP mouse antibody (1:200) in the *ΔMyAP::MyAPΔEFh-Ty* strain (green). ConA lectin surface staining of epimastigotes in red. DAPI in blue.

**F.** Immunofluorescence assay of MyAP using the αMyAP mouse antibody (1:200) in the *ΔMyoC* strain (green). ConA lectin surface staining of epimastigotes in red. DAPI in blue.

**G.** Immunofluorescence assay of MyAP using the αMyAP mouse antibody (1:200) in the *ΔMyoF* strain (green). ConA lectin surface staining of epimastigotes in red. DAPI in blue.

**H.** Immunofluorescence assay of MyAP using the αMyAP mouse antibody (1:200) in the *ΔMyoF::ΔMyoC* strain (green). ConA lectin surface staining of epimastigotes in red. DAPI in blue.

Scale bars 2 μm.

**Supplementary Figure 7 Endocytosis is Not Disrupted by the Mutagenesis of Predicted Calcium Binding Residues in MyAP**

**A.** AlphaFold structure showing three aspartic acid residues that we identified as potentially essential for calcium binding in the EF-hand structure, with D616 being the most promising candidate.

**B.** Expanded gel (from Figure 5 Panel B) shows PCR amplification of the MyAP locus and successful heterozygous complementation of the aspartic acid to alanine mutagenized versions of MyAP (*blue dotted rectangle*) into the original locus (*faint high-MW band*).

**C.** Expanded immunoblot (from Figure 5 Panel C) probed with αMyAP shows successful expression of the mutagenized aspartic acid versions of MyAP (*blue dotted rectangle*).

**D.** Quantification of three fluorescent BSA feeding assays shows that all three mutagenized versions of MyAP (*yellow, orange, dark orange*) were capable of fully rescuing endocytic activity in the *ΔMyAP* background.

**Supplementary Figure 8 Principal Component Analysis of Mass Spectrometry-Based Lipidomics Analysis**

**A.** The sample type (media versus *T. cruzi*) is separated along principal component 1 (PC1). The separation along PC2 indicates that the Parental and Complement share similarities with the Media, whereas the KO is distinct from those three. Principal-component analysis of lipidomic data reveals that *T. cruzi* strains lacking the ability to endocytose (*ΔMyAP*) display an alteration of the lipid profile of parasites grown in LIT/LDNT (Media) as compared to those strains able to carry out SPC mediated nutrient uptake (Parental and *ΔMyAP::MyAP-Ty* complemented lines) as highlighted in the blue dotted line rectangle.

## Notes

### Competing Interest Statement

The authors have declared no competing interest.

### Summary of Updates

Inclusion of Supplementary Figures

