## Supplementary figures and images for "Endocytosis in *Trypanosoma cruzi* Depends on Proper Recruitment and Regulation of Functionally Redundant Myosin Motors"

### Supplemental Figure 1

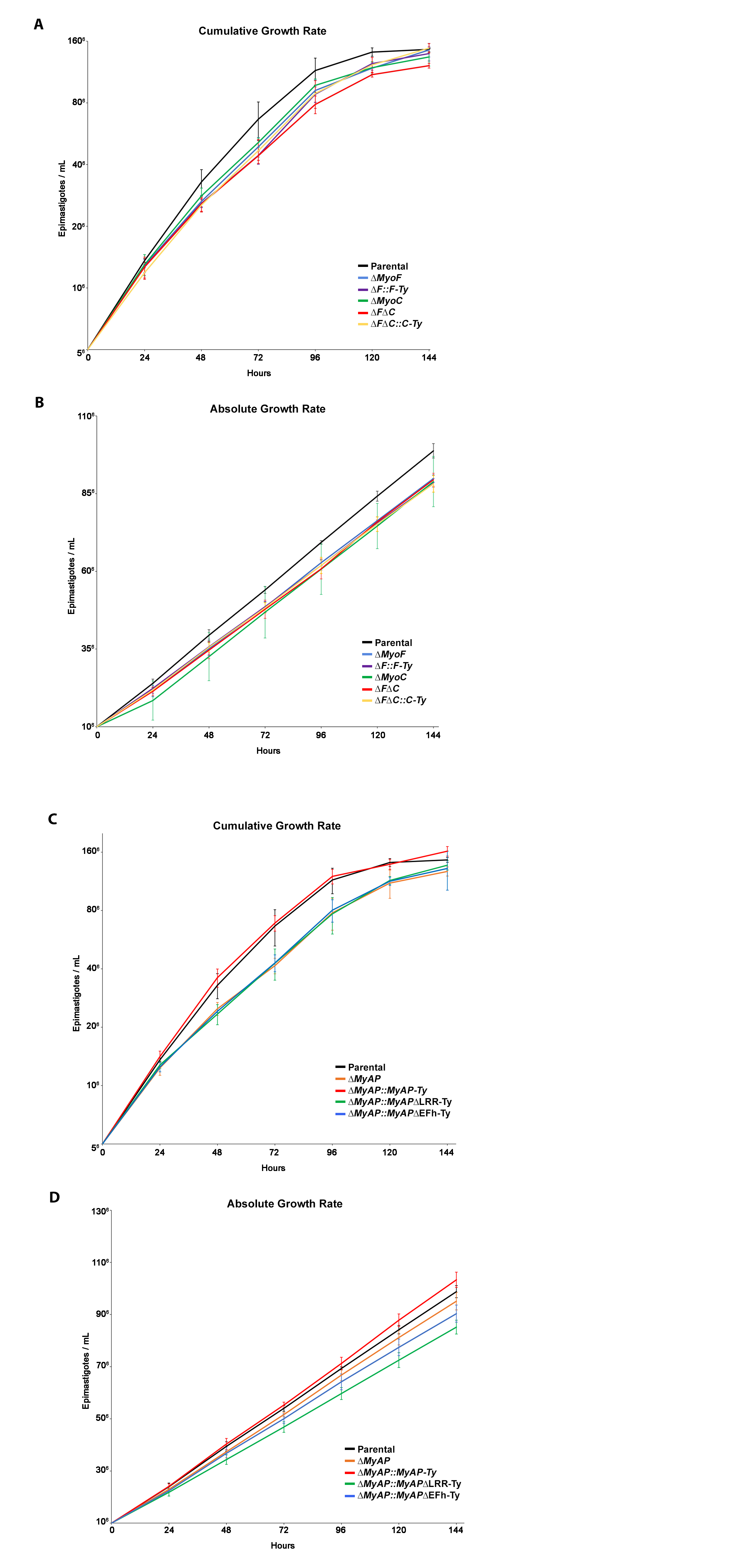

### Supplemental Figure 2

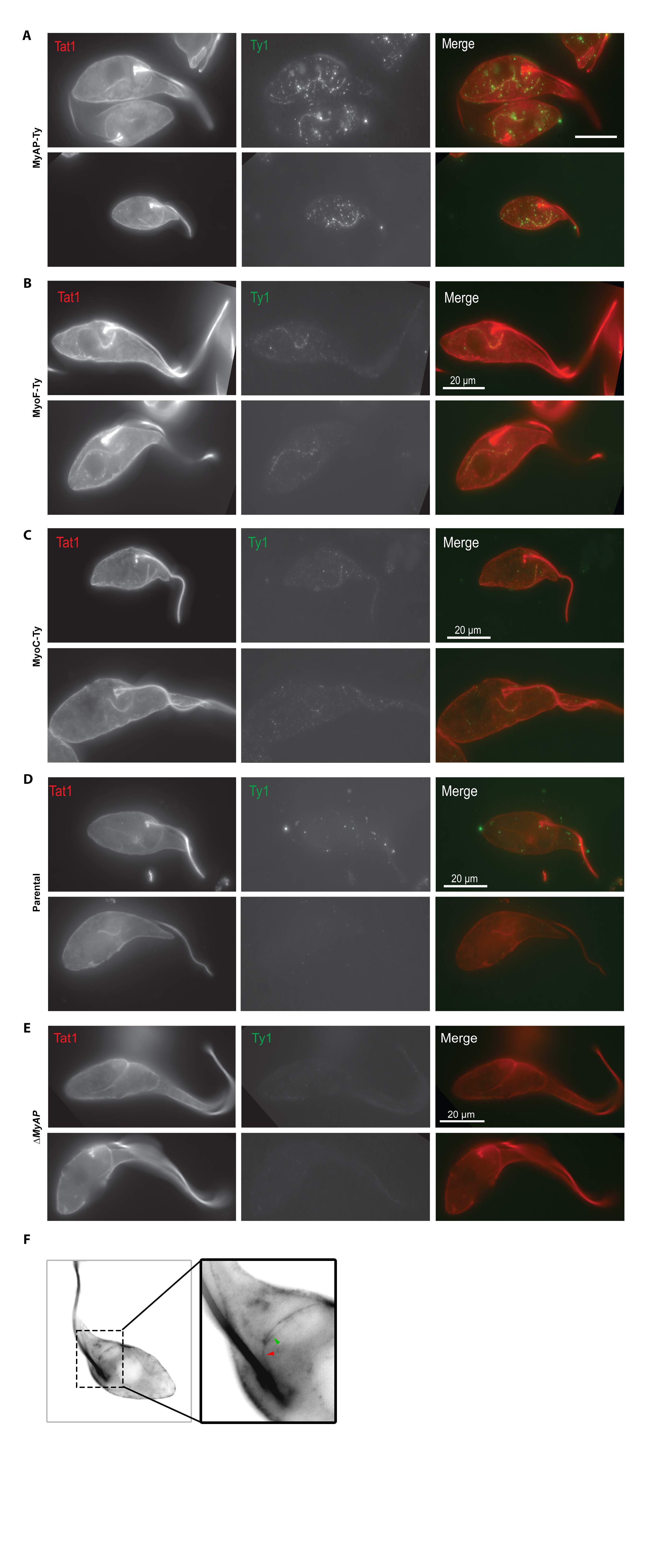

### Supplemental Figure 3

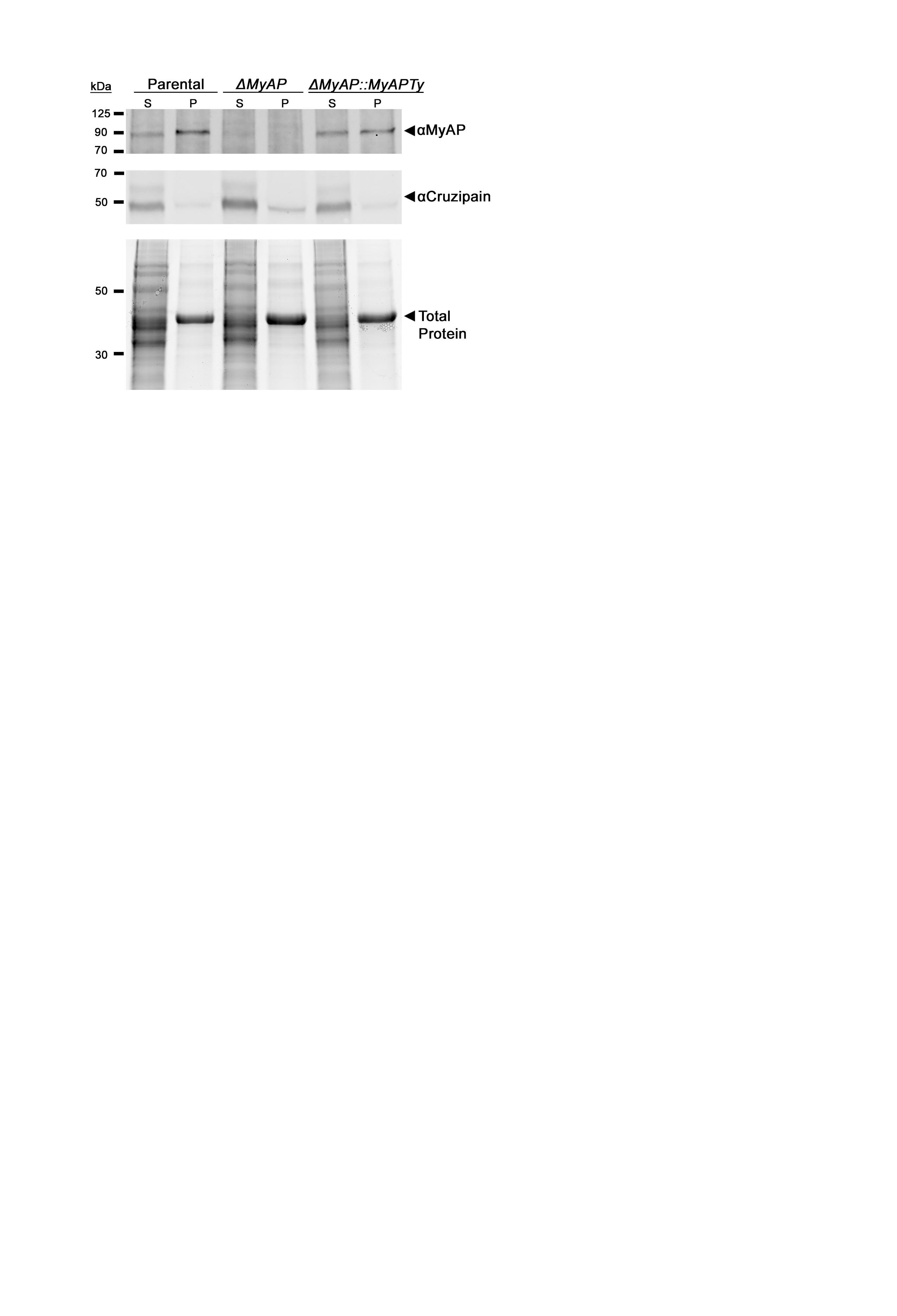

### Supplemental Figure 4

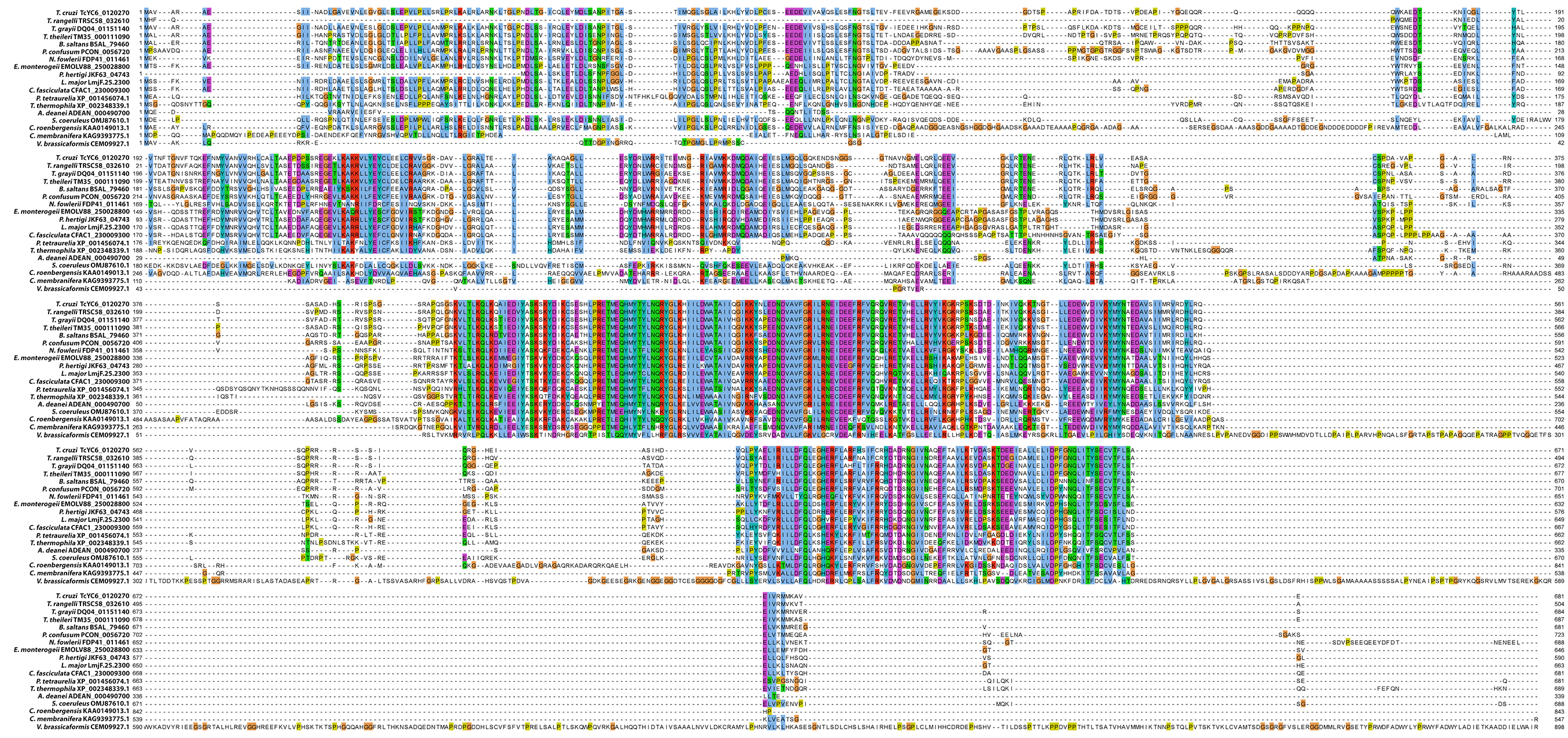

### Supplemental Figure 5

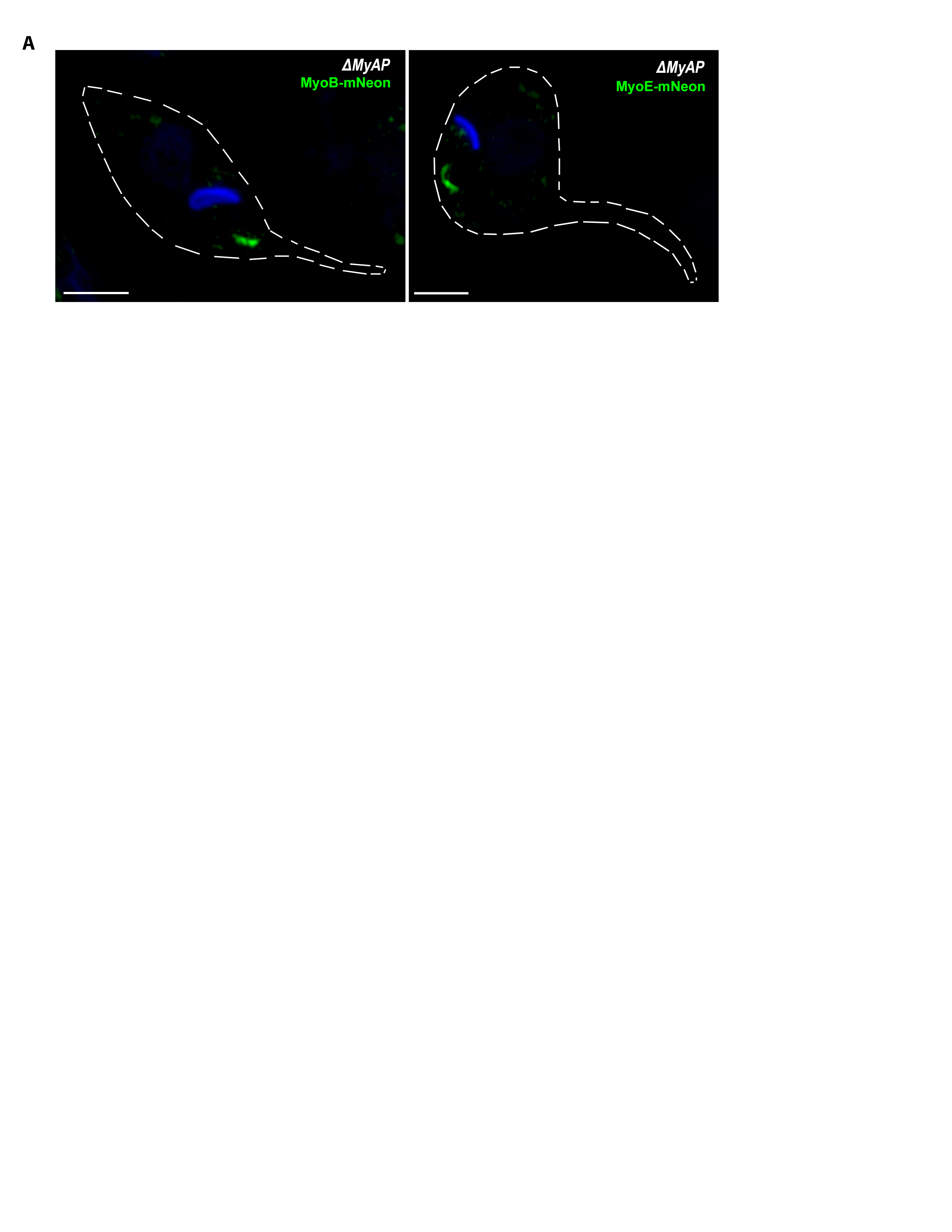

### Supplemental Figure 6

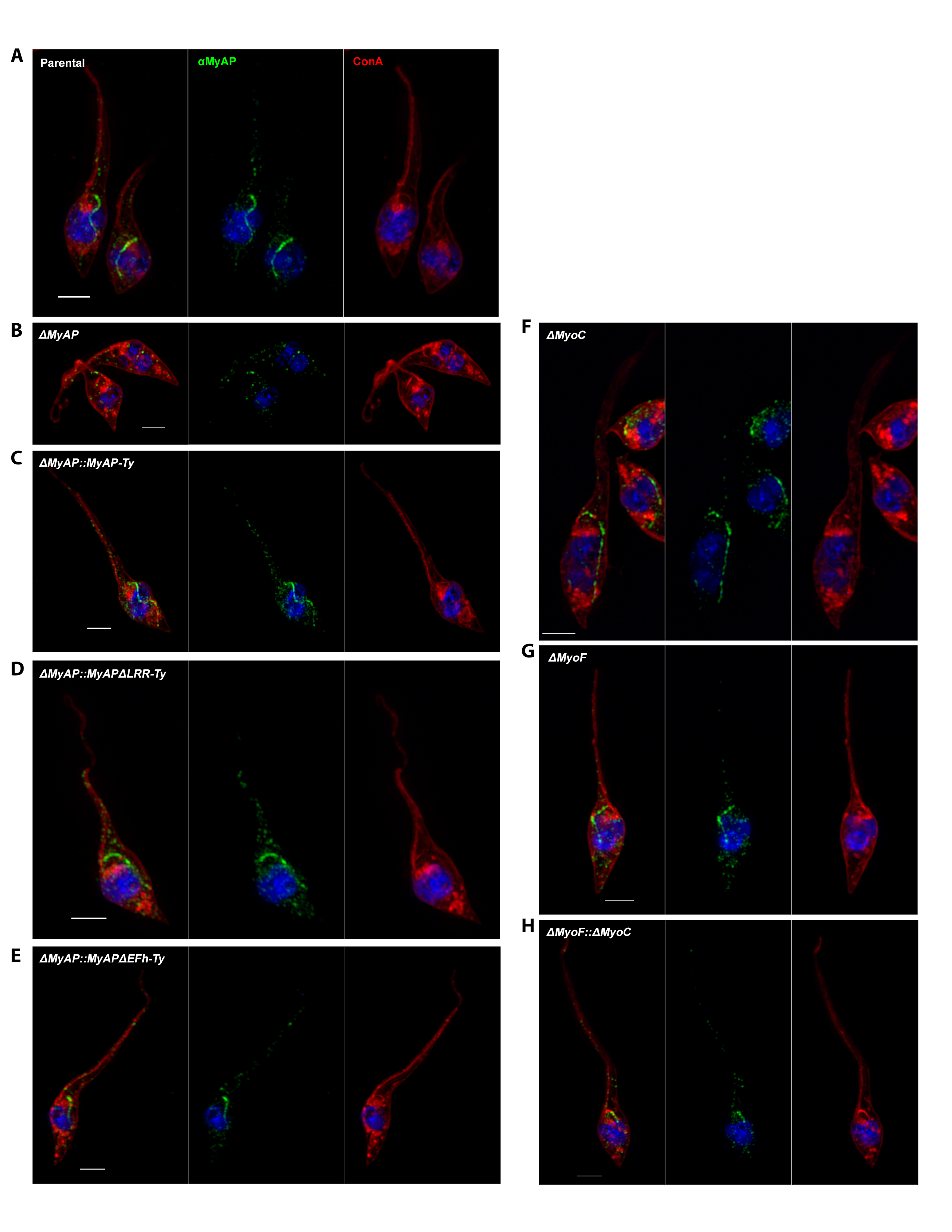

### Supplemental Figure 7

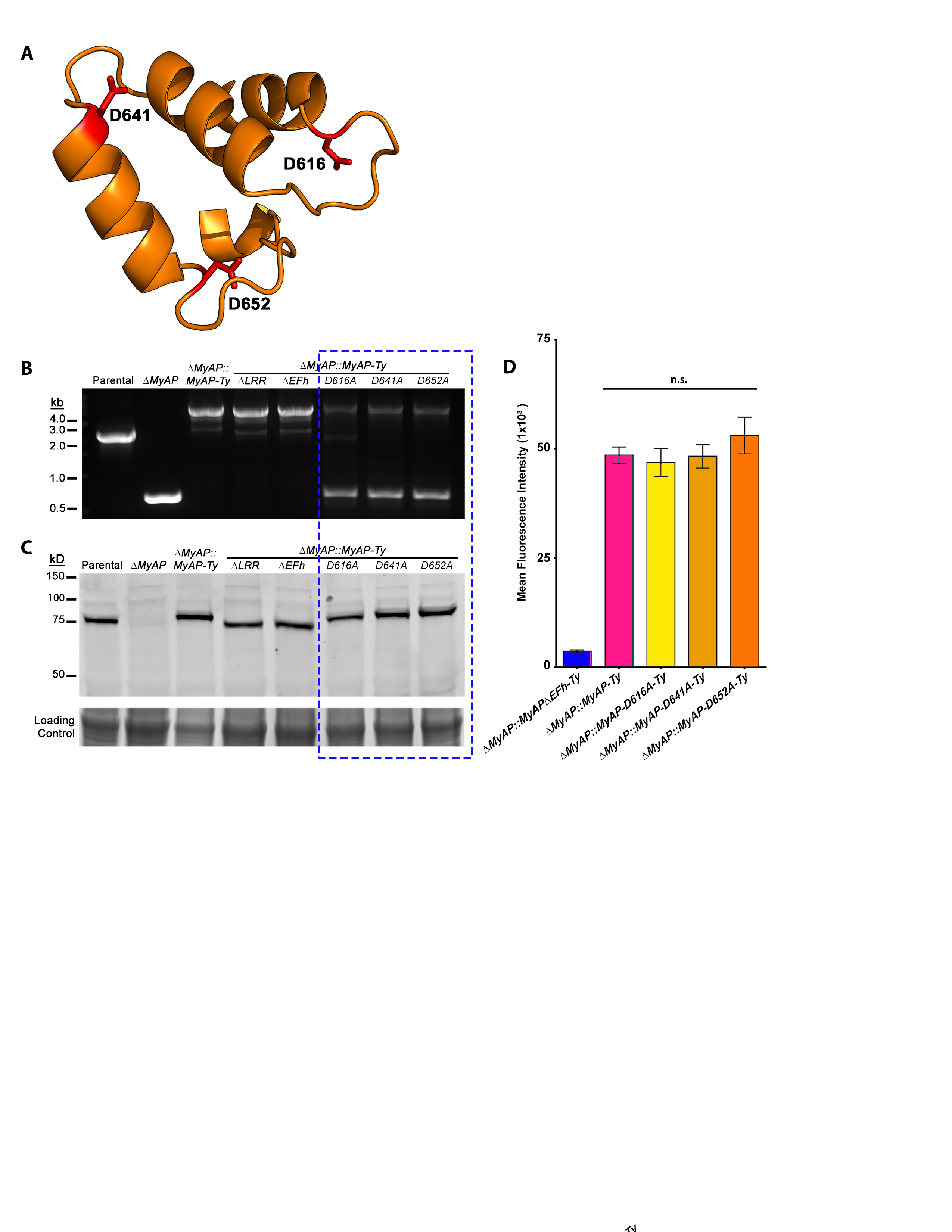

### Supplemental Figure 8

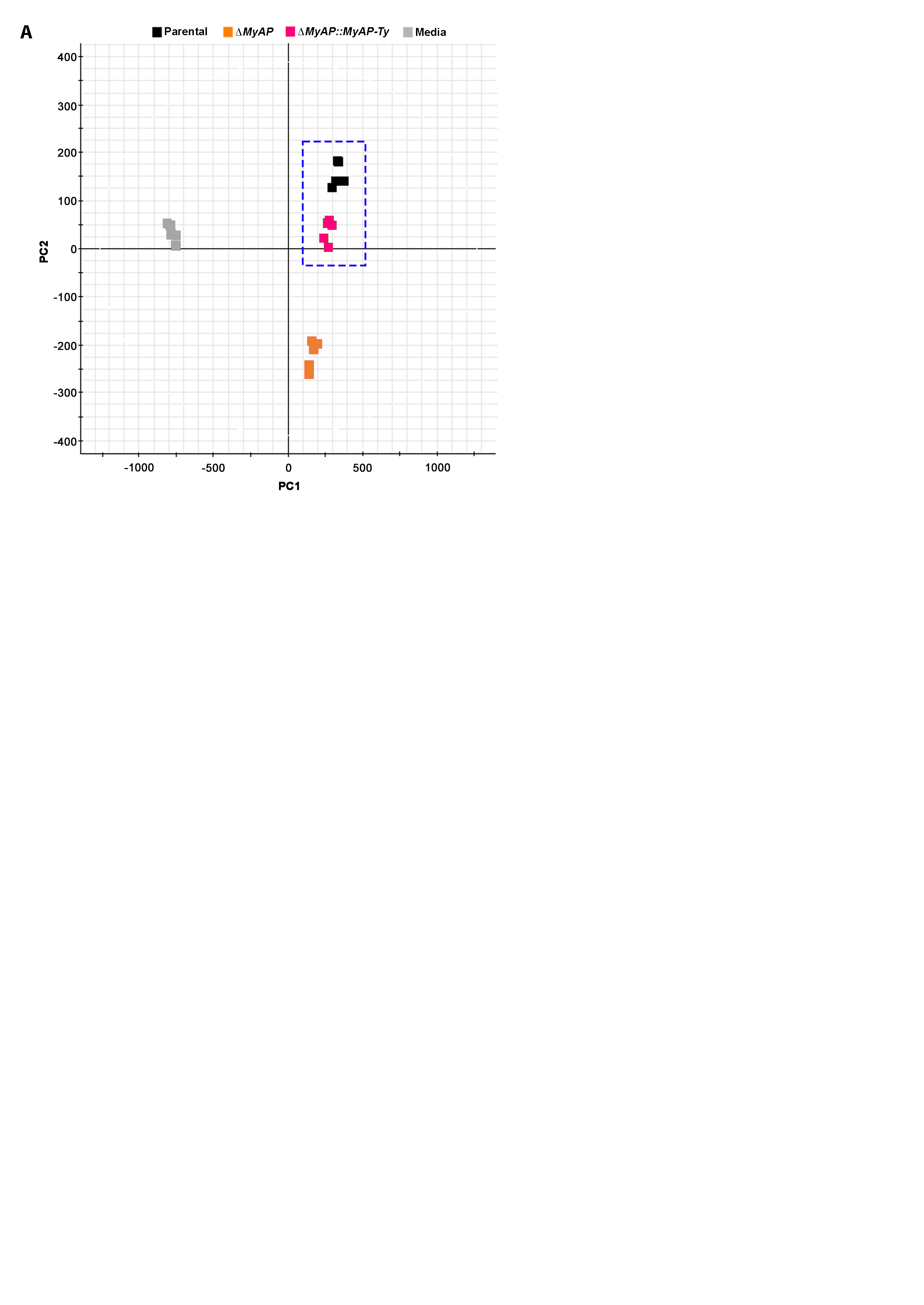
